# Distinct Type 1 Immune Networks Underlie the Severity of Restrictive Lung Disease after COVID-19

**DOI:** 10.1101/2024.04.03.587929

**Authors:** Glenda Canderan, Lyndsey M. Muehling, Alexandra Kadl, Shay Ladd, Catherine Bonham, Claire E. Cross, Sierra M. Lima, Xihui Yin, Jeffrey M. Sturek, Jeffrey M. Wilson, Behnam Keshavarz, Naomi Bryant, Deborah D. Murphy, In Su Cheon, Coleen A. McNamara, Jie Sun, Paul J. Utz, Sepideh Dolatshahi, Jonathan M. Irish, Judith A. Woodfolk

## Abstract

The variable etiology of persistent breathlessness after COVID-19 have confounded efforts to decipher the immunopathology of lung sequelae. Here, we analyzed hundreds of cellular and molecular features in the context of discrete pulmonary phenotypes to define the systemic immune landscape of post-COVID lung disease. Cluster analysis of lung physiology measures highlighted two phenotypes of restrictive lung disease that differed by their impaired diffusion and severity of fibrosis. Machine learning revealed marked CCR5+CD95+ CD8+ T-cell perturbations in mild-to-moderate lung disease, but attenuated T-cell responses hallmarked by elevated CXCL13 in more severe disease. Distinct sets of cells, mediators, and autoantibodies distinguished each restrictive phenotype, and differed from those of patients without significant lung involvement. These differences were reflected in divergent T-cell-based type 1 networks according to severity of lung disease. Our findings, which provide an immunological basis for active lung injury versus advanced disease after COVID-19, might offer new targets for treatment.

## Introduction

Persistent complications following SARS-CoV-2 infection are well-recognized, and include persistent cough, dyspnea, hypoxemia, and interstitial lung disease^1–10^. Risk is especially high among patients who were hospitalized with severe COVID illness, and symptoms may persist for months or years after infection^2,8^. However, the underlying immune mechanisms remain enigmatic owing to the complex clinical landscape of post-acute sequelae of COVID-19 ^2,11^, the heterogeneous nature of lung disease, and lack of comprehensive lung assessments in parallel with immune measures.

T cells are pivotal to chronic inflammatory disorders of the lungs, including those that may culminate in fibrosis.^12^ Although T-cell responses to SARS-CoV-2 infection have been extensively documented during acute illness^13,14^, the nature of T cells involved in the pathophysiology of post-acute disease is less clear. Survivors of severe COVID-19 display alterations in numbers of naïve, effector, and virus-specific CD4+ and CD8+ T-cell subsets in the blood, as well as polarization in their functional states^15–19^. Moreover, patients with persistent post-acute symptoms display dysregulation of T cells and other inflammatory mediators in the blood that imply an ongoing immune response.^20^ Nonetheless, the relationship between peripheral T cells and lung damage is unclear, perhaps due to localization of pathogenic T cells to the airways and/or their very low numbers in the blood. While T-cell immunophenotyping strategies in blood and airways have begun to tackle this problem^16,21^, there is a critical need to better understand immune mechanisms underlying poor pulmonary outcomes.

In this study, we applied a novel multi-modal analytical strategy to define T-cell immune networks in the blood that promote lung damage after COVID-19 illness. This was done by leveraging a longitudinal cohort of 110 COVID-19 survivors that is unique for its systematic assessment of lung physiology and immune measures spanning up to 20 months after infection. By coupling unbiased machine learning and integrative methods, we identify discrete immune networks underlying the severity of restrictive lung disease after COVID-19. Our ability to discriminate the systemic immune landscape of active lung injury from that of more advanced disease might inform pharmacological targets to halt disease progression.

## Results

Patients (n=110) primarily presenting to a post-COVID clinic with dyspnea on exertion (DOE), cough, and/or fatigue were enrolled into a COVID Recovery Cohort (COVID-RC)(**Figs. 1a-b**). Most had been hospitalized for severe illness (84%) and were on mechanical ventilation (62%)(**Fig. 1b; Supplementary Fig. 1**). The majority of patients became ill before the emergence of viral variants and vaccine availability (**Supplementary Fig. 2**). Patients received a comprehensive evaluation of the respiratory system, and returning patients (n=31) were longitudinally monitored up to 20 months post-COVID (**Supplementary Fig. 1**). Blood specimens were also collected for comprehensive immune profiling. Comparison groups used to establish COVID-specificity consisted of patients with mild COVID-19 sampled at acute infection and 1 month, as well as uninfected healthy and vaccinated controls, and subjects with obesity (**Fig. 1a**).

**Figure 1.**
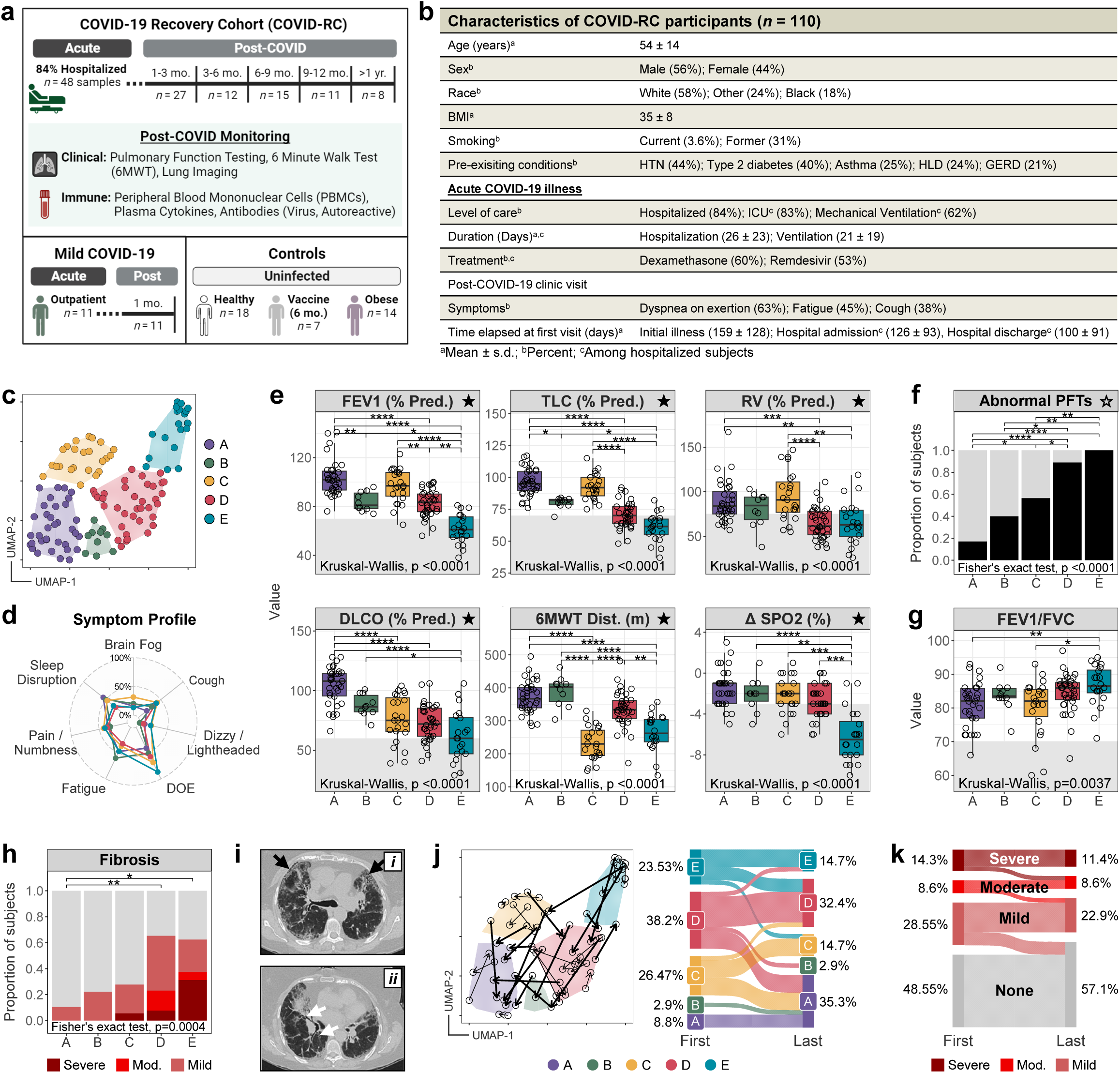
Clinical Phenotypes of Pulmonary Disease in a COVID-19 Recovery Cohort. **(a)** Study schematic for the COVID-19 Recovery Cohort (COVID-RC, *n* = 110 patients), including comparison groups (Mild COVID-19, *n* = 11; Healthy, *n* = 18; Vaccine, *n* = 7; Obese, *n* = 14). Sample sizes in figure correspond to the number of specimens used in cellular studies. **(b)** Characteristics of COVID-RC patients. **(c–k)** Unsupervised analysis of lung physiology measures (*n* = 87 subjects, 124 visits). **(c)** Visualization of 5 patient groups (A: *n* = 35, B: *n* = 10, C: *n* = 23, D: *n* = 36, E: *n* = 20). **(d)** Percent of subjects reporting symptoms at follow-up visits. **(e)** Representative lung physiology data by patient group. Shading denotes abnormal values. **(f)** Proportion of subjects with any abnormal PFT. **(g)** FEV1/FVC ratio by cluster. **(h)** Proportion of patients with fibrosis (*n* = 65 subjects, 88 visits). Statistics reflect fibrosis versus no fibrosis. **(i)** Representative chest CT scans of severe fibrosis (9 months after COVID-19 positive test), with extensive associated peripheral honeycomb change (***i***, black arrows), and extensive associated peripheral traction bronchiectasis and bronchiolectasis (***ii***, white arrows) comprising more than 50% of total lung volume, including all five lung lobes. **(j)** Transitions between clusters over time (*left*; *n* = 34 subjects, 71 visits), and from first to last visit (*right*; *n* = 34 subjects). **(k)** Change in fibrosis from first to last visit in the whole COVID-RC (*n* = 35 subjects). ★, measures used to derive clinical phenotypes; ☆, categorizations also derived from ★ measures. 6MWT, 6-minute walk test; Dist., distance; Mod., moderate. Kruskal-Wallis with Dunn’s post-hoc test and Holm adjustment (***e, g***), and Fisher’s exact test (***f, h***). \**P* ≤ 0.05; \*\**P* ≤ 0.01; \*\*\**P* ≤ 0.001; \*\*\*\**P* ≤ 0.0001.

### Cluster Analysis Identifies Distinct Pulmonary Phenotypes

To first address clinical heterogeneity, post-COVID patients with complete lung measure datasets (104 subjects, 124 visits) were classified according to pulmonary function measures (spirometry, lung volume, and diffusing capacity of the lungs for carbon monoxide [DLCO]) and 6-minute walk test ([6MWT], distance and oxygen saturation). Data dimensionality reduction with cluster analysis (UMAP with self-organizing maps^22,23^) yielded five patient clusters denoted A through E (**Fig. 1c**). The highest prevalence of patient-reported DOE was observed in C through E, and the severity of lung disease increased from A to E according to objective lung measures (**Figs. 1d-f; Extended Data Figs. 1a-e**). Most patients in groups D and E, but not C, had restrictive lung disease based on lung volume measures (total lung volume [TLC] and residual volume [RV]; 72% and 90% versus 4%), whereas obstructive airway disease (decreased FEV1/FVC) was not prominent in any cluster (<15%)(**Figs. 1e,g; Extended Data Figs. 1a-c**). Moreover, inspection of available chest CT scans (65 patients, 88 visits) identified more than 60% of patients with evidence of interstitial fibrosis in groups D and E, with the greatest prevalence of severe fibrosis in E (**Figs. 1h-i, Supplementary Fig. 3**).^24^ Notably, group E was also characterized by the most impaired gas exchange (DLCO), and differed from all groups by marked desaturation after walk test (**Fig. 1e; Extended Data Figs. 1b,d-e**).

Despite a lack of airway restriction, patients in group C demonstrated the worst mobility by both objective (6MWT) and subjective metrics (self-reported mobility impairment, ability to perform daily activities, pain/discomfort), but were not discriminated by desaturation after walk (**Fig. 1e; Extended Data Figs. 1a-f**). Instead, ∼40% had reduced DLCO and ∼17% displayed air trapping (**Extended Data Figs. 1b-c**). This could be explained by abnormal breathing patterns and autonomic dysfunction after COVID-19, often diagnosed as chronic fatigue syndrome/myalgic encephalitis, postural orthostatic tachycardia syndrome and post-exertional malaise (hereafter termed “other functional impairments”).^25–27^ Accordingly, a few individuals had decreased heart rate responses and abnormal reduction in blood pressure after exercise (**Extended Data Figs. 1g-h**). By contrast, and in keeping with their reduced DOE, groups A and B were defined by limited pulmonary impairment and elevated diastolic blood pressure (B only)(**Figs. 1e-g; Extended Data Figs. 1g**).

Analysis of transitions between phenotypes for sequential visits (n=34) found that 53% of patients remained in the same clinical cluster, 41% tracked improvement in the direction of E to A, while a minority (6%) moved in the opposite direction (**Fig. 1j**). In contrast, only 5 of 39 patients (13%) with repeat lung imaging had improved fibrosis scores, indicating persistent lung changes up to 794 days post-discharge (**Fig. 1k**). Time since illness did not vary by patient group (**Extended Data Figs. 2a**). Review of the clinical record for relationships to risk factors associated with fibrosis (Age, BMI, sex, race, smoking, comorbidities), SARS-CoV-2 vaccination and antibodies, and acute clinical care revealed only trends toward male (B and D) or female (A and C) predominance by group, and increased ventilator days in E when excluding non-ventilated patients from analysis (**Extended Data Figs. 2b-h; Supplementary Table 1**). Together, these findings demonstrated that groups D and E comprised distinct phenotypes of restrictive disease, with E constituting the most severe fibrosis, gas exchange impairment and desaturation, consistent with more advanced disease.

### Profound CD8+ T-Cell Dysregulation Delineates Restrictive Lung Phenotypes

To understand the immunopathogenesis of lung sequelae, we next asked whether different phenotypes of restrictive lung disease linked to distinct T-cell signatures in the blood. T-cell deep phenotyping was performed by spectral flow cytometry in the COVID-RC (48 acute and 73 convalescent samples, 55 subjects) and comparison groups (Mild COVID-19; Healthy, vaccinated, and obese controls; 61 samples, 50 subjects). The resulting data was dimensionality reduced (**Figs. 2a-b, Extended Data Figs. 3a-c**), and the machine learning tool Tracking Responders Expanding (T-REX)^28^ was applied to pinpoint disease-relevant populations, including rare and novel cell types, based on their degree of change in comparison with health (≥95% increase or decrease)(**Supplementary Fig. 4**). This strategy revealed marked T-cell dysregulation in COVID-RC patients when compared with mild COVID-19, and changes persisted beyond one year (**Extended Data Figs. 3d-e**). Cell alterations impacted multiple CD4+, CD8+, and γδ T-cell populations, and patterns were distinct for each pulmonary phenotype (**Figs. 2a-d**). Notably, T-cell perturbations were most marked for restrictive phenotype D, and comparatively attenuated in phenotype E (**Figs. 2d-e**). T-cell changes in group D were dominated by enrichment of two “metaclusters” (denoted Mc1 and Mc2). Each of these contained similar cell populations based on Root-Mean-Square Deviation (RMSD) analysis of complex phenotypic signatures identified by Marker Enrichment Modeling (MEM)^29^ (**Figs. 2d-e; Supplementary Table 2**). Populations in both Mc1 and Mc2 expressed the type 1 transcription factor T-bet, the lung-homing marker CCR5, and CD95 (Fas), but differed by their activation and differentiation states (**Figs. 2f-g**). Mc1 contained mostly activated (HLA-DR+) effector memory cells (T_EM_, CD45RA-CCR7-) spanning CD45RO+ non-proliferating and CD45RO^lo^CD45RA+ proliferating (Ki67+) types (**Fig. 3a-c**; **Supplementary Fig. 6**). Interestingly, the activation marker CD38 was not a feature. In contrast, Mc2 contained TEMRA-like cells (CD45RA+CCR7-CD27-) spanning non-activated non-proliferating and activated (HLA-DR+CD38+) proliferating types (**Figs. 3g-i**), consistent with pro-inflammatory and cytotoxic terminally-differentiated cells found in blood.^30–34^ Expansion of Mc1 and Mc2 during acute illness, their elevation compared with mild COVID, and their persistence and expansion thereafter, indicated their robust induction during initial infection (**Extended Data Figs. 4a-b**). Importantly, these populations were not induced in uninfected obese or vaccinated controls, suggesting that expansion was infection-specific (**Supplementary Fig. 5**, *not shown*). Interestingly, three TEMRA-like populations correlated with improved lung measures (**Figs. 3j**).

**Figure 2.**
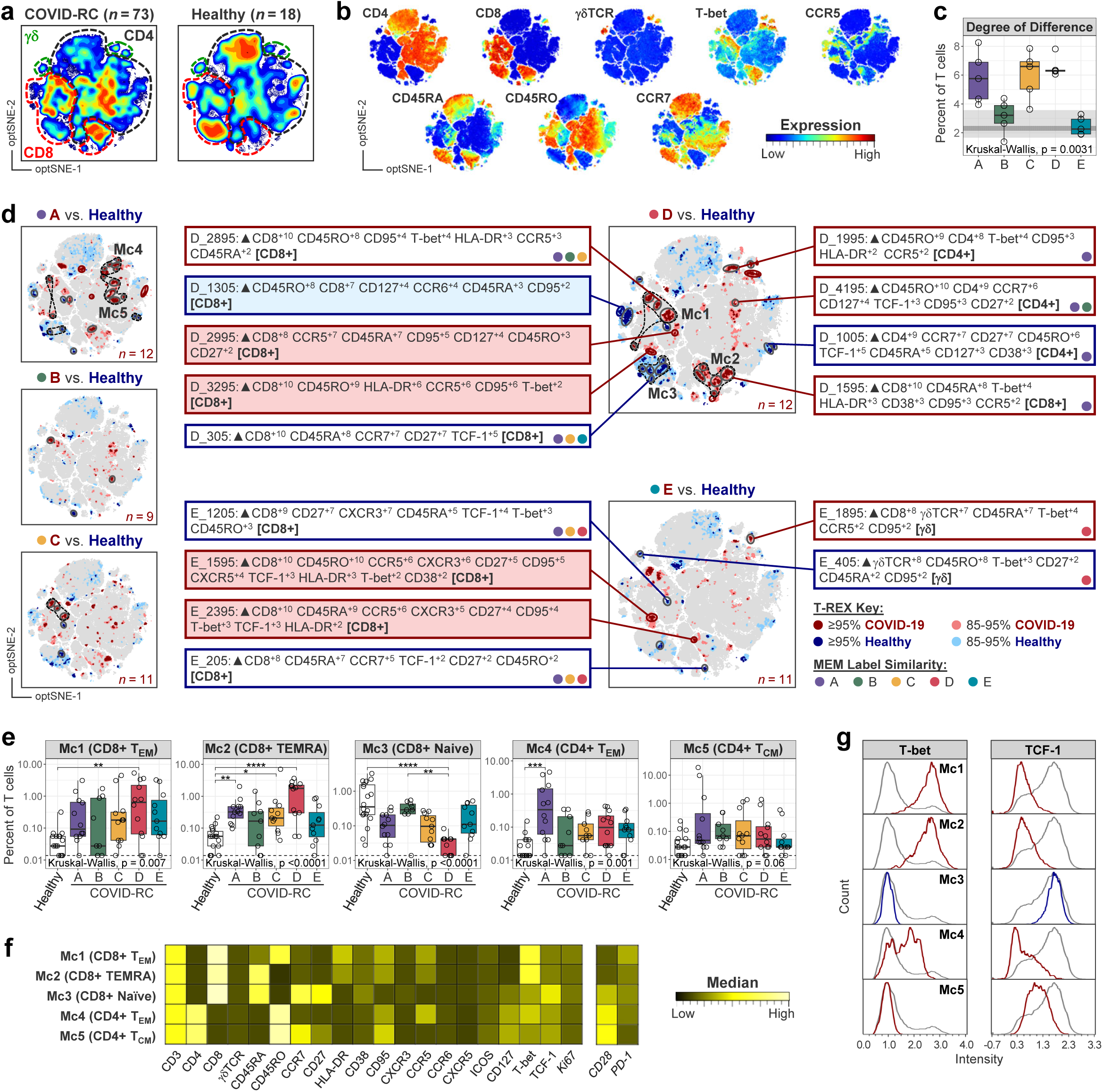
Distinct T-Cell Populations Link to Pulmonary Phenotypes. **(a)** opt-SNE dimensionality reduction of CD3+ T cells in the COVID-RC and healthy groups. CD4+, CD8+, and γδT cells are annotated. **(b)** Select marker expression plots. **(c)** T-REX Degree of Difference (percent of cells in regions of 95% enrichment) in COVID-RC versus healthy controls, by patient group. Analyses iterated excluding single batches (*n* = 5). Shading shows the range and IQR of healthy controls (see **Supplementary Fig. 4**). **(d)** T-REX plots for each patient group, as compared with healthy controls (*n* = 18). Red populations are increased in COVID; blue populations are increased in healthy (decreased in COVID). Validated robust populations are circled (red/blue, unique to each patient group; gray, shared), and representative MEM labels are shown for D and E (red/blue shading, unique; no shading, shared). Dashed outlines denote metaclusters (Mc) 1-5, and colored dots in MEM label boxes indicate signature similarity across patient groups, as determined by RMSD (cutoff, 90). T-cell type is annotated in brackets. **(e)** Frequency of metaclusters according to pulmonary phenotype. **(f)** Comparison of median marker expression between metaclusters in the COVID-RC (samples with low cell counts are excluded: *n* ≥ 62; PD-1 & CD28, *n* ≥ 56). Markers in italics were not used in opt-SNE analysis. **(g)** Transcription factor profiles of metaclusters. Concatenated data, with total T cells shown in gray. Kruskal-Wallis with Dunn’s post-hoc test and Holm adjustment (***c, e***). \**P* ≤ 0.05; \*\**P* ≤ 0.01; \*\*\**P* ≤ 0.001; \*\*\*\**P* ≤ 0.0001.

**Figure 3.**
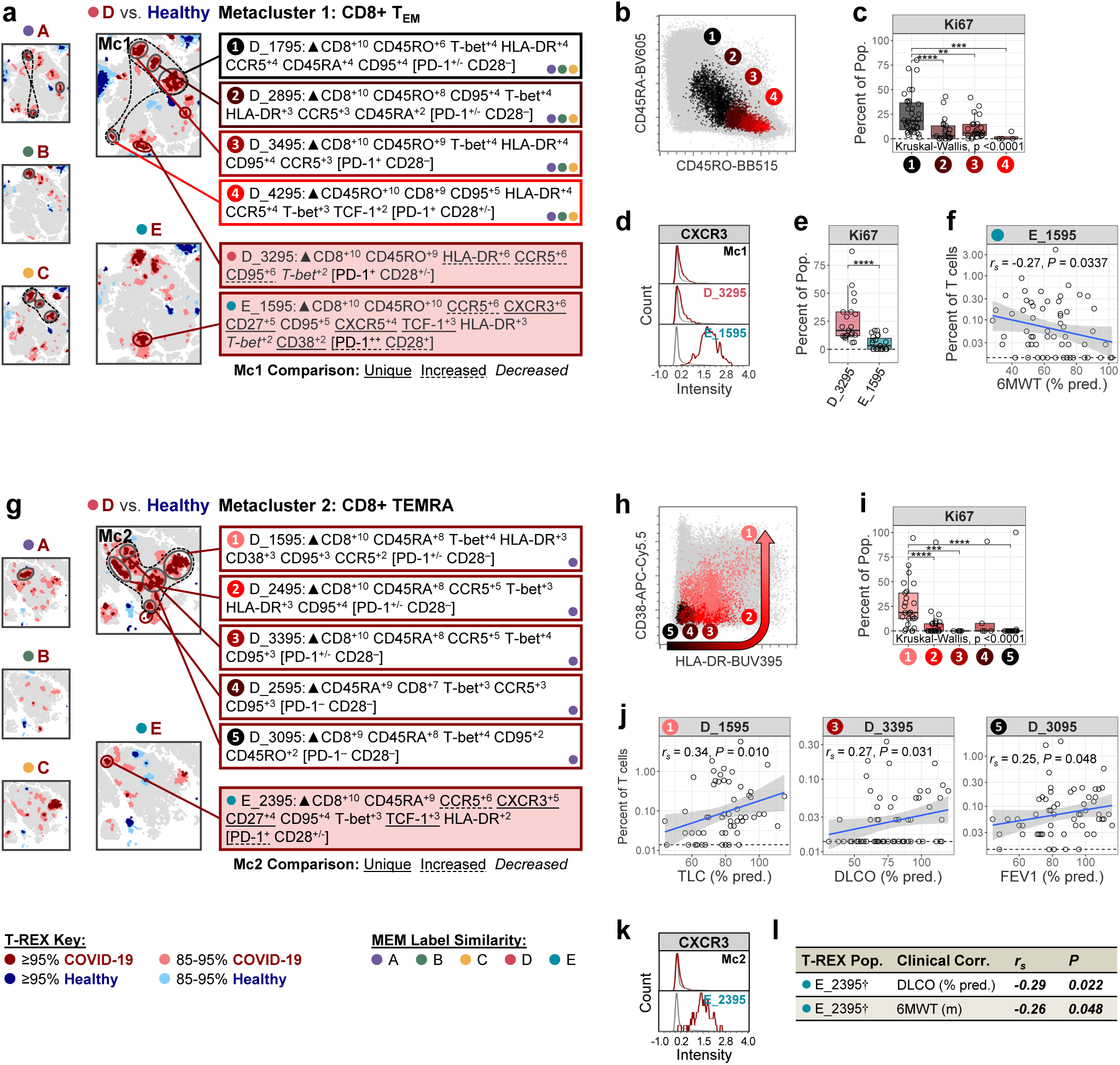
Features of CD8+ T-cell Populations Enriched in Patients with Restrictive Lung Disease. **(a)** CD8+ T_EM_ populations in Mc1 with corresponding MEM labels, as identified by T-REX analysis of group D (*n* = 12) vs. healthy controls (*n* = 18) and RMSD. Adjacent populations unique to D (D_3295) or E (E_1595) are shown in shaded boxes, with distinguishing features highlighted. **(b)** Expression of CD45RA and CD45RO across Mc1. **(c)** Cell proliferation (Ki67) across Mc1 (*n* = 86; ≥4 per pop.). **(d)** Expression of CXCR3 in CD8+ T_EM_ populations unique to groups D and E. **(e)** Cell proliferation in unique T_EM_ populations (*n* = 44; ≥21 per pop). **(f)** Spearman correlation between unique population E_1595 frequencies and 6MWT data in the COVID-RC (*n* = 60). **(g)** CD8+ TEMRA populations in Mc2 with corresponding MEM labels, as identified by T-REX analysis of group D (*n* = 12) vs. healthy controls (*n* = 18) and RMSD. An adjacent population unique to E (E_2395) is shown in a shaded box, with distinguishing features highlighted. **(h)** Expression of CD38 and HLA-DR (arrow denotes increasing activation) across Mc2. **(i)** Cell proliferation across Mc2 (*n* = 92; ≥5 per pop). **(j)** Significant spearman correlations between Mc2 frequencies and lung function measures in the COVID-RC (*n* ≥57). **(k)** Expression of CXCR3 in a CD8+ TEMRA population unique to group E. **(l)** Significant spearman correlations between unique population E_2395 and lung function measures in the COVID-RC (*n* = 60). E_2395 contains ≥70% zero frequencies (†). PD-1 and CD28 (measured in 4 of 5 batches) were excluded from T-REX analyses and MEM labels, and expression is summarized in brackets (***a, g***). Colored dots in MEM label boxes denote signature similarity across patient groups, using an RMSD cutoff of 90 (***a, g***). Concatenated data, total T cells shown in gray (***b, d, h, k***). Samples with low cell counts within each population were excluded from analysis (***c, e, i***). Kruskal-Wallis with Dunn’s post-hoc test and Holm adjustment (***c, e, i***). \**P* ≤ 0.05; \*\**P* ≤ 0.01; \*\*\**P* ≤ 0.001; \*\*\*\**P* ≤ 0.0001.

Restrictive phenotypes D and E also differed by the enrichment of CCR5+CD95+CD8+ T-cell populations that were unique to each group (**Figs. 3a,g**). Those unique to E (E_1595, E_2395) contrasted with D (D_3295) based on expression of CXCR3, CD27, and TCF-1; higher PD-1 and lower Ki-67; and included CXCR5+PD-1^hi^ T_EM_-like (E_1595), and CD45RA+CCR7-CD27+ (E_2395) signatures (**Figs. 3a,d-g,k; Supplementary Fig. 6**). Although rare, their frequencies inversely correlated with lung function (DLCO, 6MWT)(**Figs. 3f,l**) supporting a pathogenic role for unique T-cell signatures in patients with advanced lung disease.

In addition to enriched CD8+ T-cell metaclusters, patients in group D displayed marked depletion of a third metacluster (Mc3) (**Figs. 2d-e**) of naïve CD8+ T-cells displaying varying expression of CXCR3 and CD38 (**Extended Data Figs. 4c-d; Supplementary Fig. 6**), including “effector-like” CXCR3+ cells (D_2105, D_705).^35^ A marked decrease during acute illness, persistent depletion during convalescence in group D, and comparatively modest decreases in obese individuals (**Extended Data Figs. 4e-f**) likely reflect sustained recruitment of naïve T cells to the CD8 T-cell response in inflammation, including in patients with restrictive lung disease.

### CD4+ T-Cell Populations Link to Improved Lung Function

Patients with restrictive lung disease frequently transitioned from group D to A (**Fig. 1j**), consistent with recovery of lung function. Seeking commonalities in T-cell populations between these groups that might contribute to lung disease resolution identified scarce Mc1- and Mc2-like populations in group A (**Fig. 2d**). However, the T-cell landscape was dominated by two CD4+ T-cell metaclusters comprising CCR5+ T_EM_ (Mc4) and T_CM_ (Mc5) populations (**Figs. 2d-e; Extended Data Figs. 4g**). In contrast, similar CCR5+ CD4+ T_EM_ identified in group D (D_1995, D_2795) expressed higher T-bet and HLA-DR, and were more proliferative (**Figs. 2f-g; Extended Data Figs. 4h-j**, **Supplementary Fig. 6**), consistent with increased activation in lung disease. CD4+ T_CM_ (Mc5) correlated with N-specific IgG antibodies, and both Mc4 and Mc5 populations correlated with improved lung function (**Extended Data Figs. 4l**). Moreover, while CD4+ T_CM_ were expanded during acute illness, they were markedly contracted in convalescence (**Extended Data Figs. 4m-n**), in contrast with the dynamics of potentially pathogenic CD8+ T-cell responses. Together, these observations implied protective functions of Mc4 and Mc5.

### Candidate Pathogenic and Protective T Cells are Present in the Airways of COVID Patients

We next asked whether T-cell signatures identified in the blood after COVID-19 were present in the lungs of patients who had prior COVID-19 using leftover specimens from clinically indicated bronchoscopies. CCR5, a key feature of these signatures, was ubiquitously expressed in T-cells in the airways (**Extended Data Figs. 5a-c**). Three populations corresponding to candidate T-cell signatures were enriched in bronchoalveolar lavage (BAL) versus blood: (1) BAL_14, similar to Mc1 (CD8+ T_EM_); (2) BAL_12, similar to unique population E_1595 (CXCR3+CXCR5+PD-1^hi^ CD8+ T_EM_); and (3) BAL_40, similar to Mc4 (CD4+ T_EM_)(**Extended Data Figs. 5d-e**). Mc2-like (CD8+ TEMRA) cells were also identified in the BAL of some patients, but were not enriched (*not shown*). Notably, CXCR3 was a feature of all enriched populations, as were markers of tissue residency (CD69, CD103)(**Extended Data Figs. 5d,f**). These data demonstrate that candidate T cells are positioned to exert pathogenic or protective functions in the airways.

### Virus-Specific T Cells are Elevated in Patients with Restrictive Lung Disease

Activation-Induced Marker (AIM) assays to analyze frequencies of virus-specific T cells in COVID-RC patients found increased Spike (S)- and Nucleocapsid (N)-specific CD4+ and CD8+ T cells in restrictive phenotypes D, and E, as well as C. Positive correlations with secreted IFN-γ, IL-2, and TNF-α indicated functionality (**Extended Data Figs. 6a-b**). Comparing the *ex vivo* T-cell profiles of patients with the highest (AIM^hi^) and lowest (AIM^lo^) numbers of virus-specific T cells using T-REX revealed enriched CD8+ Mc-1-like cells in AIM^hi^ individuals (**Extended Data Figs. 6c**). Further analysis revealed that Mc1 frequencies positively correlated with N-specific CD8+ T cells (**Extended Data Figs. 6d**). Notably, Mc1 populations were the only populations that were enriched in group C according to T-REX (**Fig. 2d**). By contrast, CD4+ Mc4- and Mc5-like cells were depleted in AIM^hi^ patients, and frequencies of Mc4 cells and virus-specific CD4+ T cells were negatively correlated (**Extended Data Figs. 6c-d**). Together, these data suggest that virus-specific CCR5+CD95+CD8+ T_EM_ cells contribute to persistent T-cell dysregulation in patients who have restrictive lung disease or other functional impairments (group C), and that virus-specific CD4+ T cells are likely non-protective from post-acute sequelae in these patients.

### Distinct Immune Networks Underlie the Severity of Lung Disease

Contrasting T-cell profiles in patients with restrictive lung disease pointed to divergent inflammatory states underlying mild-to-moderate (group D) and advanced disease (group E). To better understand these differences, comprehensive immune profiling was performed, including measurement of plasma cytokines and chemokines (76 mediators) and autoantibodies targeting immune mediators and traditional autoantigens (73 targets)(**Extended Data Figs. 7a-b**). Broad immune monitoring of cellular responses was performed in tandem by manual gating of canonical immune cell subsets (110 populations) including both T- and non-T populations (**Extended Data Figs. 7c**). Inspection of these datasets identified elevated CXCL13 as the sole cytokine discriminator of group E from all other groups. Group E was also notable for having the highest levels of type 1 and pro-inflammatory mediators (CXCL10, TNF-α) and the chemokine CCL21 (**Extended Data Figs. 7a**). To further harness the complexity of these data, we used a multi-modal integrative method. Elastic net regularization and orthogonalized partial least squares-discriminant analysis (OPLS-DA) were applied to define a minimal set of immune features that discriminated each patient group from healthy controls (**Fig. 4a**).^36–38^ For rigor, this method excluded T-REX populations (defined by contrasting COVID-19 and health) and data from AIM assays (sample size). As this approach eliminates co-linear features that might also discriminate groups, co-correlates (*R* ≥0.8) were also interrogated (**Extended Data Figs. 8**).

**Figure 4.**
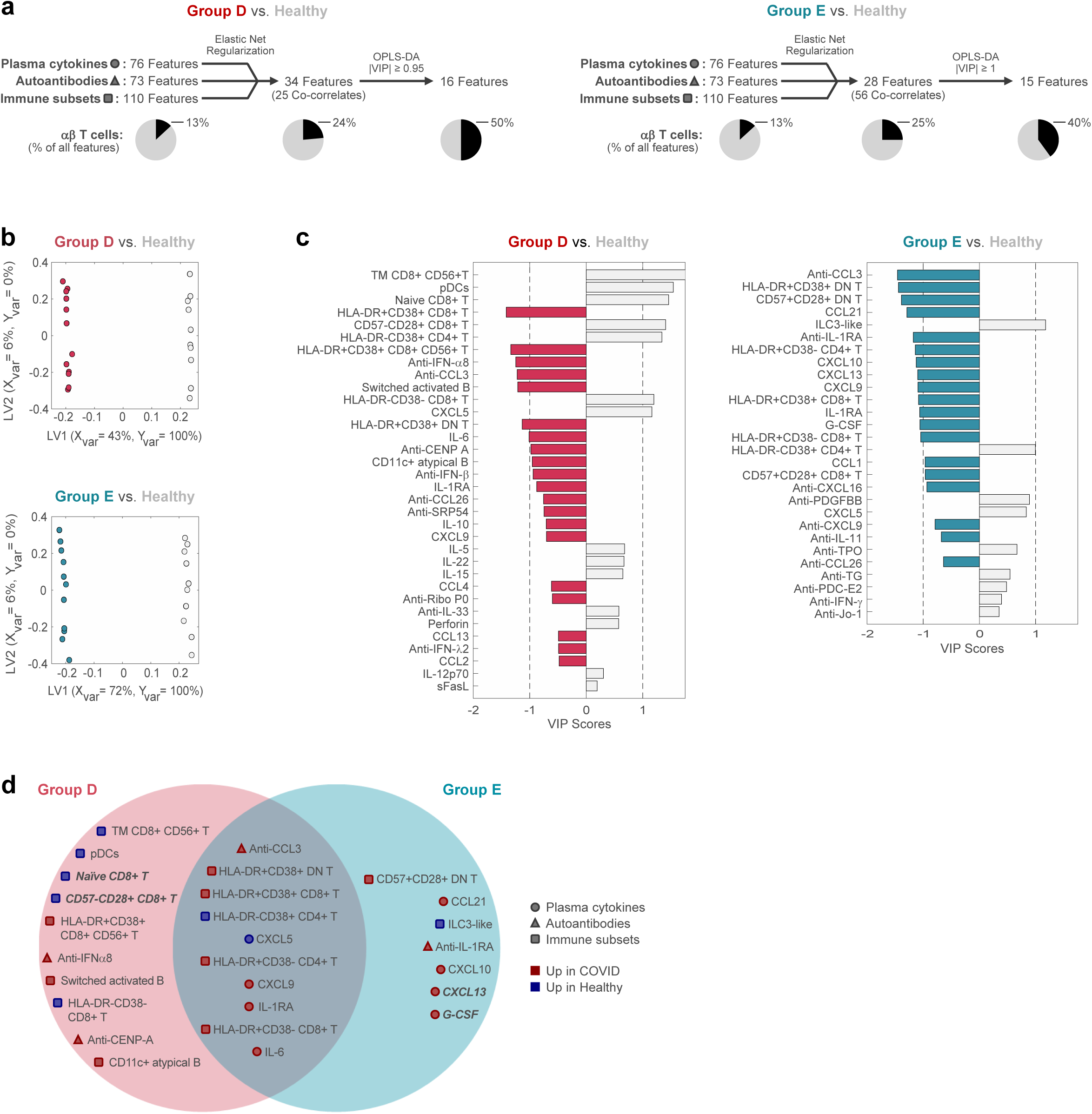
Distinct Immune Profiles Discriminate Restrictive Lung Disease Phenotypes D and E. OPLS-DA models were created using elastic net-selected features that discriminated group D or E from healthy controls. **(a)** Integrative analysis pipeline for groups D (*left*) and E (*right*). Elastic net feature selection was run separately for each dataset before performing combined OPLS-DA. The percent of features that correspond to αβ T cells (CD3+γδTCR-) are denoted. **(b)** OPLS-DA score plots depicting models of separation for groups D (*top*; *n* = 12) or E (*bottom*; *n* = 11) versus healthy controls (*n* = 10), where each point represents one sample. Use of an orthogonalized approach ensured that latent variable 1 (LV1) captures between-group separation while simultaneously capturing 100% of the Y-variation, while LV2 captured variances that did not contribute to this separation. 5-fold cross-validation (CV) resulted in 100% CV accuracy for both models. Both models significantly outperformed models based on shuffled group labels (permutation testing, Wilcoxon *p*=0.002 for both). **(c)** Ordered Variable Importance in Projection (VIP) scores of all elastic net-selected features for the discrimination of group D (*left*) or E (*right*) from healthy controls. **(d)** Comparison of the top discriminants of groups D or E versus healthy (|VIP| ≥0.95 or 1, respectively), ordered according to VIP score. Features are shared if they also appear as a discriminant (no VIP cutoff) or a co-linear feature (**Extended Data Figs. 9**) for the alternate group. Symbols denote data type (shape), and whether features are increased in post-COVID patients or healthy controls (color). Features that also discriminate D vs. E in a direct elastic net comparison are indicated in bold.

Distinct sets of immune measures provided perfect separation of groups D and E from healthy controls (**Fig. 4b**). T-cell features were among the top contributors to both models (|VIP score| ≥0.95 or 1), and these manually gated populations were consistent with those previously identified by T-REX, including increases in HLA-DR+CD8+ T cells (D and E) and reductions in naive CD8+ T cells (D). Other notable shared discriminants and co-correlates included increased levels of CCR5 and CXCR3 ligands (CCL4 and CXCL9), autoantibodies targeting CCR5 ligands (anti-CCL3), as well as inflammatory cytokines (IL-6) and immunomodulatory factors (IL-1RA)(**Fig. 4c-d**). Features unique to group D included B-cell populations (switched activated and CD11c+ atypical). By contrast, those unique to group E included chemoattractants and growth factors linked to lung fibrosis (CXCL13, G-CSF, CXCL10) (**Fig. 4c-d**). Type 2 and IL-17 family members were also co-correlates of group E discriminants, in keeping with a mixed inflammatory milieu (**Extended Data Figs. 8b**).

To elucidate immune networks in groups D and E, including the contributions of T-REX populations and data from AIM assays, correlation networks were generated for discriminants and co-correlates within each patient group. Resulting networks were distinct, and shared discriminants correlated with different sets of immune features within each group (**Supplementary Table 3**). For example, while the shared discriminant CXCL9 negatively correlated with non-activated CD8+ T cells in both groups D and E, all other correlations with immune cell subsets were distinct (8 in D, 14 in E)(**Extended Data Figs. 9a**). To examine candidate pathogenic T-cell populations in the context of the larger immune landscape, we next explored networks centered around T-REX populations. Within group D, T-REX Mc1 and Mc2 populations (CD8+ T_EM_ and TEMRA) and the unique population D_3295 (CD8+ T_EM_) positively correlated with pro-inflammatory cytokines, autoantibodies, and manually gated cell subsets, including activated (HLA-DR+CD38+/-) CD8+ T cells (**Fig. 5a**). Non-activated non-proliferative TEMRAs (D_3095, Mc1) did not positively correlate with any feature and negatively correlated with cytokine responses, consistent with hyporesponsiveness. Group D networks also prominently featured autoantibody discriminants including anti-IFNs, and were notable for inclusion of B-cell subsets including atypical B cells (implicated in autoimmunity) that correlated with a CD4+ T_CM_ population in Mc5 (A_2995) within a pathogenic context. Within group E, unique CXCR3+ T-REX populations (E_1595, E_2395) positively correlated with activated T cells (**Fig. 5b**). Of note, the TEMRA-like population E_2395 negatively correlated with the pro-fibrotic factors CXCL13 and CCL21. Given that CXCL13 was the sole discriminant of group E according to both univariate and multivariate methods, we next explored the complete correlation network surrounding this mediator (**Fig. 5c**). CXCL13 positively correlated with 17 cytokines and chemokines, including type 2-associated, IL-17 family, and pro-fibrotic mediators, thrombopoietin (THPO), as well as non-specific (PHA) and N-specific cytokine secretion in AIM assays and gdT-cell populations.

**Figure 5.**
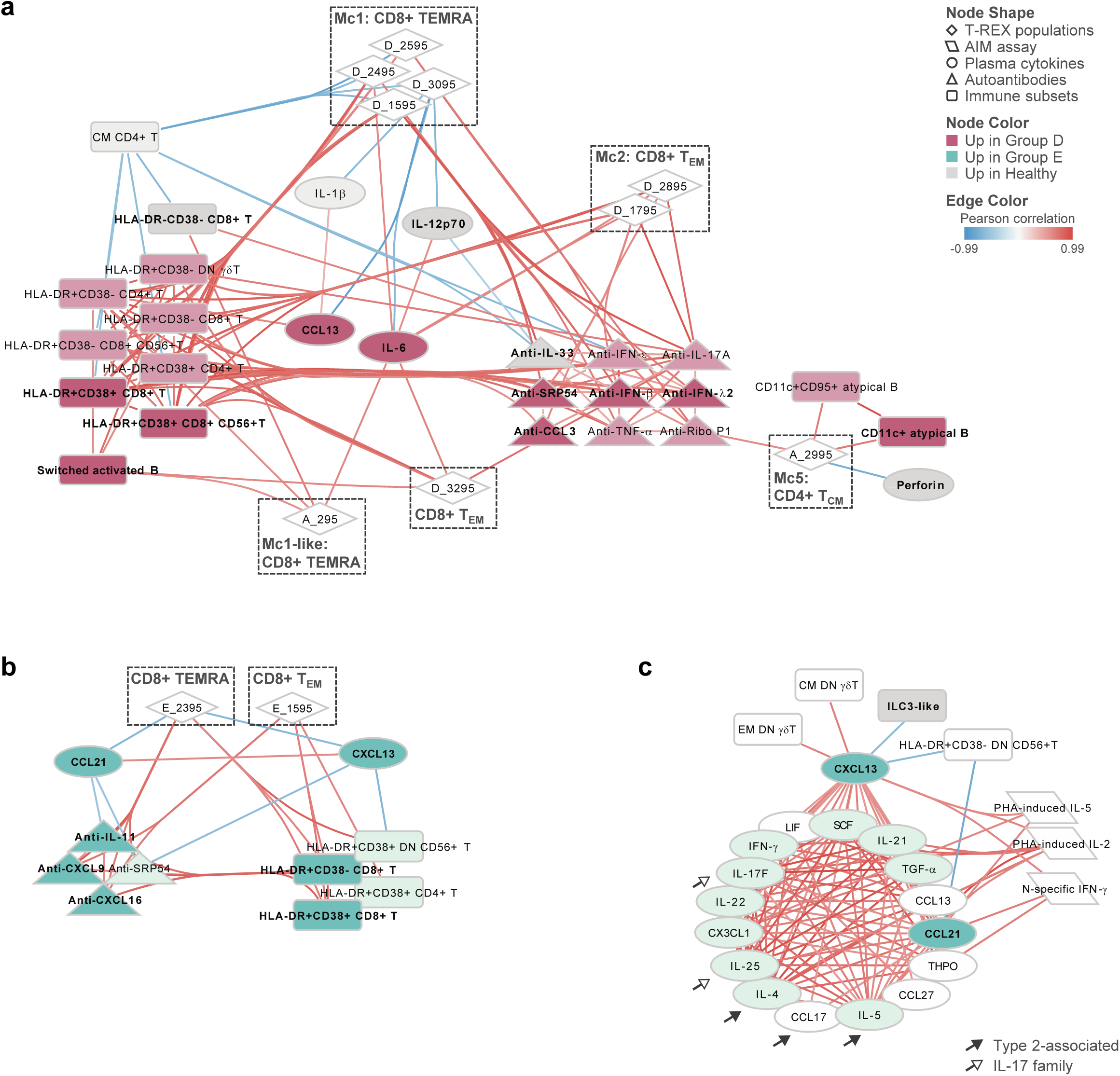
Immune Networks That Underlie Restrictive Airway Disease Phenotypes D and E. **(a & b)** Pearson correlation networks generated within **(a)** group D (n ≥ 11), and **(b)** group E (n ≥ 11). Networks depict all features that significantly correlate with T-REX populations identified within each group (denoted with dashed boxes). Group D networks (***a***) also include T-REX populations identified in group A for comparison. **(c)** Pearson correlation network generated within group E (n ≥ 11), depicting features that significantly correlate with CXCL13. For readability, correlations of CXCL13 with T-REX populations, autoantibodies, and immune subsets that were previously depicted in ***b*** are not shown here. Type 2-associated and IL-17 family mediators are highlighted by arrows. For all, data utilized in correlation analyses include T-REX population frequencies, AIM assay data (cell frequencies and secreted cytokines), and SARS-CoV-2-specific IgG, in addition to data utilized in elastic net and OPLS-DA modeling (plasma cytokines, autoantibodies, immune subset frequencies). Node shape denotes data type, and color denotes elastic net-selected features (dark) and co-correlates (light); edge color represents Pearson *R* (*P_adj_* ≤0.05).

Immune networks were next compared with non-restrictive phenotypes, including those with other functional impairments (group C) or largely normal lung function (A). Discrimination of group C from healthy controls was less robust than for D and E, potentially due to underlying disease heterogeneity (**Extended Data Figs. 10a-b**). Although group C shared some discriminants with D and E, and Mc1-like T-REX populations (C_1295, C_2095) also correlated with activated CD8+ T cells, fewer features discriminated group C as compared with restrictive phenotypes (**Extended Data Figs. 10c-e**). In contrast to E, anti-CXCL13 was a unique discriminant of group C, and has been suggested to protect against post-COVID sequelae.^39^ The network for group A was distinct from all other groups, and notable for the inclusion of CD4+ CD56+ T-cell discriminants that correlated with Mc4 and Mc5 populations (**Extended Data Figs. 10f-j**). In contrast to group D, no significant correlations between CD4+ T_CM_ populations (Mc5) and atypical B cells were observed within group A, emphasizing the distinct immune mechanisms of pathogenic versus protective responses. Accordingly, networks around shared features revealed altered relationships within advanced lung disease (E) versus groups A and C, including inverted correlations between the shared chemokine CCL21 and switched activated B cells or diverse γδT-cell populations (E—negative; A and/or C—positive)(**Extended Data Figs. 9b**). These collective findings highlight distinct immune programs that distinguish restrictive lung disease phenotypes from each other, and from other post-acute phenotypes of disease.

## Discussion

Persistent breathlessness, one of the most common symptoms after COVID-19 illness, can emanate from disease in various organ systems.^2,40^ Consequently, the complex etiology of post-acute sequelae has hindered efforts to delineate the immunopathology of lung complications. Here, we leveraged a comprehensive set of lung physiology measures in COVID patients to identify two discrete phenotypes of restrictive lung disease that encompassed milder and more advanced forms of disease. A key strength of our approach was the ability to not only discriminate the severity of fibrotic lung disease using lung physiology measures alone, but to separate patients with lung disease from those with other disorders typical of “long COVID” diagnoses. Further, by longitudinal tracking, we were able to identify transitions consistent with improvement in lung disease. Our strategy for rigorous categorization of patients was foundational to deciphering a distinct immunological basis for different types of restrictive lung disease after COVID-19. Use of complementary single-cell machine learning and multi-modal methods demonstrated a link between vigorous type 1 dysregulation consistent with active lung injury in patients with mild-to-moderate restrictive lung disease. By contrast, more advanced disease was characterized by a comparatively attenuated T-cell landscape and a mixed inflammatory profile hallmarked by CXCL13.

In patients with mild-to-moderate disease, the T-cell landscape was dominated by expanded T-bet+ CD8+ T_EM_ and TEMRA-like cells that expressed CCR5 and CD95. Both types were armed for lung homing and displayed signatures consistent with recent activation in tissue.^41,42^ CCR5 is rapidly induced on T cells during acute respiratory viral infection, is critical to early migration of memory CD8+ T cells to the lungs^43,44^, and together with its ligands (CCL3, CCL4, CCL5) has been implicated in the pathogenesis of interstitial lung diseases and acute COVID-19 illness.^43–49^ In accord, CCL4, along with CXCL9, another type 1 chemoattractant, was integral to inflammatory signatures of patients in group D. Although CXCR3, the receptor for CXCL9, was not a prominent feature in the blood, low levels were expressed on T cells present in the lungs of COVID patients (Mc1, CD8+ T_EM_), suggesting its upregulation in lung tissue where it may contribute to tissue repair or else promote virus-induced lung pathology.^50,51^ CXCR3 was also expressed on “effector-like” naïve CD8+ T-cells that were chronically depleted in the blood, most likely reflecting their egress out of circulation for participation in the T-cell response.^35^ These collective data, together with transitional states of activation, differentiation, and proliferation within CD8+ T_EM_ and TEMRA metaclusters, implied a highly dynamic T-cell response whereby protracted activation sustains a cytotoxic, pro-inflammatory TEMRA pool.^31,52–54^ These expanded CD8+ T-cell populations are key players in a robust immune response characterized by B cells, pro-inflammatory cytokines, and anti-interferon antibodies.

The immune features of patients with more advanced lung disease (group E) were strikingly different to those of group D. T-cell dysregulation in the blood was attenuated, even in the face of a mixed inflammatory milieu comprising type 1, type 2, and IL-17 family mediators, and growth factors involved in fibrosis. Moreover, elevated plasma CXCL13 was a unique hallmark in this group, building on its prior description as a biomarker of severity and mortality during acute COVID-19, its induction by hypoxia, and links to progressive fibrotic lung disorders.^55–58^ In our cohort, CXCL13 was induced during acute infection, and secreted by T cells upon stimulation in the chronic phase (*not shown*). Partnership between CXCL13 and CCL21, another feature of group E, may help to form and maintain inducible bronchial-associated lymphoid tissue (iBALT) during viral infection, and also aggregate T and B cells in ectopic lymphoid structures in chronically inflamed tissues.^59–63^ In accord, both CXCL13 and CCL21 are over-expressed in fibrotic foci in the lungs of COVID-19 patients where exhausted T cells accumulate.^64^ The markedly attenuated T-cell dynamics in the periphery of group E could reflect exhaustion, counter-regulation, and/or sequestration of pathogenic T cells in the lungs. Nonetheless, data from the current study suggested that rare CXCR3+PD-1+/^hi^ CD8+ T-cell populations found in the blood contributed to advanced lung disease, including a link to worse lung function.

Our results raise several important questions. First and foremost is whether the different types of restrictive lung disease we identified are related by their clinical course and immune underpinnings. Clinical transitions between groups E and D coupled with shared immune features supported commonalities. However, whether both have a similar capacity for clinical recovery requires longer term follow up. Lung disease improved in approximately one third of patients in group D, and CD4+ T cells were implicated in this process. Regarding more advanced disease, the current view is that post-COVID fibrosis is non-progressive^53,65^; however, further examination of its multi-factorial etiology, variable response to treatment^5,66^, long-term recovery, and post-acute immune drivers is warranted. The utility of CXCL13 as an early biomarker of lung disease, its role in divergent clinical courses, and potential for therapeutic targeting are all of interest in this context. While we have observed persistent immune responses following COVID, the cues that sustain this response are unknown. Hypotheses include the persistence of virus or viral antigen, and/or autoinflammation.^67^ Supporting viral persistence, we observed sustained increases in virus-specific T cells in patients with restrictive lung disease, which may contribute to pathogenic responses. Such patients might benefit from anti-viral therapies. Additionally, the link between autoreactive antibodies and atypical B cells, particulary in mild-to-moderate lung disease, pointed to autoinflammatory processes.^68–70^ Deciphering multi-factorial mechanisms of post-acute sequelae might shed light on persistent immune dysregulation, exemplified by the clinically heterogeneous patients in group C who lacked significant lung disease.

Looking forward, it will be important to probe in-depth the functions of complex T-cell populations linked to lung disease after COVID-19, as well as their cross-talk with B cells, fibroblasts, and other cells that promote fibrotic and inflammatory foci in tissue.^53,71,72^ Furthermore, understanding CD95 signaling in candidate pathogenic and protective T-cells is of interest, in light of its dual role in pro- and anti-apoptotic signaling and potential regulation of fibrosis.^73,74^ Investigating the involvement of CXCL13 in chronic pulmonary vasculopathy is also worthwhile, given its thromboinflammatory potential in COVID-19, and correlation with thrombopoietin in patients with advanced lung disease.^75–80^ Finally, validation of our findings in other cohorts will be critical, and require long-term systematic monitoring in a larger sample size. Regardless, the ability to delineate distinct systemic immune programs of lung disease constitutes an important advance in the field that could guide prognostic and therapeutic strategies for post-viral lung disease.

## MATERIALS AND METHODS

### COVID Recovery Cohort and Comparison Groups

A prospective COVID Recovery Cohort of 110 patients was initiated at the University of Virginia in June 2020. Patients enrolled into the cohort were survivors of SARS-CoV-2 infection who attended a post-COVID clinic at the University of Virginia for routine follow up care. All patients in the cohort had confirmed prior COVID-19 infection (PCR or antibody). The majority of patients were hospitalized for severe illness and most reported persistent respiratory symptoms. The cohort comprised older individuals (median age: 54 years [range 19-83 years]) and was racially and ethnically diverse (58% white, 18% black, and 30% Hispanic). Over 90% of the cohort was overweight or obese, and other medical conditions were common (**Supplementary Table 1**). 63% were unvaccinated upon enrollment, and vaccination status was documented and verified by antibody measures at each visit. Patients underwent clinical assessment at each visit, as well as pulmonary function testing with 6-minute walk test within 1 week of their visit when possible. Data was obtained from the medical record for course of illness measures during hospitalization, medications, and computed tomography (CT) scan of the lungs. Sampling of peripheral blood for cells, serum and plasma started at the initial follow up visit and continued at each visit thereafter. Matched samples collected during hospitalization were obtained through a centralized COVID Biorepository at the University of Virginia.

Patients with mild COVID-19 infection (confirmed by PCR) who did not require hospitalization (n=11; female sex, 45%; median age, 50 years [range 31-77]) were enrolled through a community screening clinic at the University of Virginia. Other comparison groups included patients who were overweight or obese (sampled before the pandemic; n=14; body mass index >29.3; female sex, 64%; median age, 52 years [range 28-64]), vaccinated healthy individuals (6 months post-vaccination; n=7; female sex, 43%; median age, 37 years [range 31-69]), and unvaccinated healthy controls who lacked antibodies to SARS-CoV-2 spike RBD (sampled from December 2020 to October 2021)(n=40; female sex, 77%; median age, 40 years [range 19-76]).

Leftover clinical bronchoalveolar lavage (BAL) and matching PBMC samples were acquired from patients with a history of COVID-19 (n=6; female sex, 50%; median age, 68 years [range 44-79]).

All studies were approved by the University of Virginia Human Investigation Committee with written informed consent (protocol numbers: 13166, 200148, 200164, 200110, 20847, 15328).

### Cluster Analysis to Define Clinical Phenotypes

Data was compiled for all clinic visits for which complete pulmonary function data was available. Dimensionality reduction was performed by UMAP (number of neighbors: 15; minimum distance: 0.01) using the following measures: percent predicted values for forced expiratory volume in 1 second (FEV1), forced vital capacity (FVC), total lung capacity (TLC), residual volume (RV), diffusing capacity of the lungs for carbon monoxide (DLCO); distance in meters and percent predicted value on 6-minute walk test; and oxygen saturation (SpO2) before and after 6-minute walk test. Data were transformed to generate all positive values. Unsupervised clustering of reduced clinical data was performed by FlowSOM (hierarchical consensus clustering; 25 clusters, 5 metaclusters, 10 iterations).^81^ All analyses were performed using Cytobank (Beckman Coulter Inc., Brea, CA).

### Assessment of Computed Tomography Scans

A pulmonologist with experience in thoracic imaging and blinded to study results reviewed all chest X-ray and CT images for the presence or absence of fibrotic pattern changes. Fibrotic pattern was defined according to the Fleischner Society glossary of terms for thoracic imaging: reticulation, architectural distortion, traction bronchiectasis, and honey combing.^82^ The severity of fibrotic pattern change was visually graded on a simplified scale modified from Li ^24^, indicating mild (1-25%), moderate (26-50%), or severe (50-99%) fibrotic pattern change as a percentage of the total visualized lung volume.

### Processing of Blood Samples for Plasma and Cells

All samples were prepared from peripheral blood using harmonized experimental protocols. 50mL blood was collected in EDTA coated vacutainers for preparation of peripheral blood mononuclear cells (PBMCs) and plasma. Briefly, blood was first centrifuged at 200g for 10 minutes to separate cells and plasma. Plasma was further centrifuged at 1000g for 10 minutes before storage at -80^0^C. PBMCs were isolated by density gradient centrifugation per manufacturer’s instructions (SepMate, STEMCELL Technologies, Vancouver, Canada) after adding DPBS 1X (Gibco, Waltham, MA) to restore initial blood volume. Cells were immediately cryopreserved in 90% FBS / 10% DMSO.

### Processing of Bronchoalveolar Lavage

Bronchoalveolar lavage (BAL) were collected for clinical indications. In brief, the bronchoscope was advanced to wedge position in the target area and three serial lavages of 50 ml of normal saline were performed. Sufficient sample volume was sent to clinical labs for diagnostic studies, and the remainder (otherwise discarded sample) was saved and processed for research. The BAL was centrifuged at 300 x g for 10 minutes. The supernatant was frozen and saved for later analysis. The cell pellet was subjected to RBC lysis (if hemorrhagic specimen), and cells were cryopreserved in 90% FBS / 10% DMSO.

### Deep Immunophenotyping of Cells

PBMCs were thawed and immunophenotyped within the same experiment using two spectral flow cytometry panels for *ex vivo* cell analysis: (1) a 24-color panel for deep immunophenotyping of T cells (panel 1), and (2) a 31-color panel designed to capture the main circulating innate and adaptive immune subsets (panel 2; described in Ayers et al^83^ (**Supplementary Table 4**). PBMC samples were run in five batches, each containing 2-3 standard batch control samples for batch normalization, and containing both patient and control samples. BAL and matching PBMC samples were run in two separate experiments and stained using spectral flow cytometry panel 1 and an overlapping T-cell panel that included tissue-resident markers (CD69 and CD103). Two million cells were stained with viability dye (LIVE/Dead Blue, Invitrogen, Carlsbad, CA, USA) and fluorescent antibodies for extracellular markers in a 100 μl mix of antibodies, Brilliant violet buffer (BD Bioscience, San Diego, CA, USA) and PBS containing 2% fetal bovine serum. Cells were then fixed and permeabilized using FOXP3 staining buffer (eBiosciences, San Diego, CA) before staining for intracellular markers. All samples were fixed before analysis to preserve signals using Cytofix buffer (BD Biosciences, San Diego, CA, USA). For optimal unmixing, reference controls were prepared using a combination of BD CompBeads Compensation Particles Set (BD Biosciences, San Diego, CA, USA) and cells from a control sample. Data for both panels were acquired by spectral flow cytometry using a 5-laser Cytek Aurora (Cytek, Fremont, CA, USA).

### Frequencies of Virus-Responsive T Cells by Activation Induced Marker Assay

T cells responding to SARS-CoV-2 proteins were measured by activation induced marker (AIM) assay.^84^ Briefly, PBMCs were stimulated in RPMI (Gibco, Waltham, MA, USA) containing 5% human serum (Sigma, Darmstadt, Germany) for 24 hours (1 million cells per well in 96-well plates) using pooled 15-mer peptides spanning the entire length of the SARS-CoV-2 original strain Spike glycoprotein and nucleocapsid (Miltenyi Biotech, Bergisch Gladbach, Germany). Cells stimulated with PHA (1 μg/ml, Roche, Basel, Switzerland) and with CEFX UltraStim peptide pool (1 μg/ml, JPT Peptide Technologies, Berlin, Germany) provided positive controls. After stimulation, cells were stained with fluorochrome-conjugated monoclonal antibodies as described in Ayers et al.^83^ (panel 3), and analyzed by spectral flow cytometry using a 5-laser Cytek Aurora (Cytek Biosciences, Fremont, CA, USA). Responding CD4+ and CD8+ T cells were identified based on OX40+CD137+ and CD69+CD137+ phenotypes, respectively. Frequencies of activated cells in non-stimulated samples were subtracted from the values in stimulated samples.

### Assay for Measurement of Cytokines in Culture Supernatants

Culture supernatants from AIM assay stored at -80^0^C were analyzed by multiplex cytometric bead assay (Milliplex, Millipore, Burlington, MA, USA) using a MAGPIX^R^ System (Luminex, Austin, TX, USA). Cytokines selected for analysis were IFN-γ, TNF-α, IL-2, IL-17A, IL-21, IL-5, IL-6, CXCL13, TGF-β1, TGF-β2, TGF-β3. Samples that did not pass quality control were excluded. For cytokines that fell outside the limit of detection, the lowest or highest value from the standard curve was assigned. For data analysis, levels of cytokines in non-stimulated samples were subtracted from the values in stimulated samples.

### Assay for Measurement of Cytokines in Plasma

Cytokines and chemokines in plasma were measured using the Human Cytokine/Chemokine 71-Plex Discovery Assay Array (Eve Technologies, Calgary, Canada) cytometric bead assay supplemented with additional analytes (Granzyme A, Granzyme B, Perforin, sFas, sFasL).

### Assay for Measurement of IgG to SARS-CoV-2 Proteins

IgG to original-strain SARS-CoV-2 spike RBD and nucleocapsid was measured in serum prepared from gold-top vacutainer tubes using a high-capacity quantitative ImmunoCAP-based assay and a Phadia 250 platform (Thermo-Fisher/Phadia, Waltham, MA, USA).^85^ In brief, SARS-CoV-2 spike RBD and nucleocapsid were biotinylated and conjugated to the streptavidin-coated solid phase of an ImmunoCAP. Unconjugated streptavidin ImmunoCAP was run in tandem with each sample and background-subtracted from the SARS-CoV-2 signal. Total IgE was also measured, as previously described.^86^

### Assay for Measurement of Autoantibodies

Plasma autoreactive IgG antibodies that target a selection of immune mediators and conventional autoimmune targets were assessed in a microbead-based antigen array assay, as previously described.^87^ Levels are reported as average MFI values, and are background subtracted for “bare bead” values. A total of 73 antigen reactivities were assessed.

### Analysis of High-Dimensional Cellular Data

All spectral flow cytometry data were pre-processed by spectral unmixing with autofluorescence subtraction and spill-over correction (SpectroFlow, Cytek Biosciences, Fremont, CA, USA). Fluorescence parameters were arcsinh-transformed with custom cofactors. Removal of dead cells, debris doublets and atypical events (antibody aggregates) was performed by expert gating. Similarly, expert gating was used to pre-gate for CD3+ lymphocytes (panel 1), and to identify known cell types/subtypes (panels 2 and 3) (FlowJo version 10.0, Tree Star Inc., Ashland, OR, USA; and OMIQ, Dotmatics, Boston, MA, USA) (**Supplementary Fig. 7**; **Supplementary Table 5**). Manual gating strategies were adapted from published works.^88–93^

#### Data Pre-Processing for T-REX Analysis

All T-cell deep phenotyping data (panel 1) was pre-gated for live CD3+ lymphocytes as previously described. Briefly, all data was dimensionality reduced using UMAP and clustered by FlowSOM in order to perform batch normalization using CytoNorm (OMIQ, Dotmatics, Boston, MA, USA). Two batch control samples were used in each experiment as internal controls; one to train batch normalization, and the other to test the validity of the normalization. Two antibodies, CD28-BV650 and PD1-BV785, were not included in batch 1, and were thus excluded from batch normalization. After normalization, MFI peaks of each fluorophore were inspected for alignment across batches, and batch control samples were inspected for the presence of non-overlapping regions between batches on dimensionality reduction maps. Between-batch T-REX analysis of batch control samples and concatenated whole batches were also performed to identify potential batch effects (**Supplementary Fig. 4**).

#### T-REX Analysis

High-dimensional data sets were analyzed by T-REX (https://github.com/cytolab/T-REX)^28^ in R (v4.2.2; RStudio, v2022.05.20-492), to resolve complex changes in cell populations over time and between subject groups. All data files were equally sampled (7,266 live CD3+ lymphocytes) prior to opt-SNE dimensionality reduction in OMIQ (perplexity, 30; theta, 0.5; iterations, 1000) (Dotmatics, Boston, MA, USA). CD28 and PD-1 were excluded from dimensionality reduction as previously described, as was the proliferation marker Ki67, to provide a confirmatory marker of recent T-cell activation in resulting populations. The T-REX modular data analysis workflow was used as previously described (**Extended Data Figs. 3a**).^28^ Briefly, data files from groups of interest were concatenated into two files for comparison, with equal sampling. K-nearest-neighbors (KNN) was employed to define an immediate phenotypic neighborhood around every cell in an opt-SNE map (k=60; population minimum cell count=50), and cell neighborhoods were assessed for the relative enrichment of neighbors from either comparison group (95% enrichment threshold). The phenotypic signatures of resulting enriched cell populations were determined by Marker Enrichment Modeling (MEM).^29^ Background marker expression was measured in known negative populations and subtracted from all MEM labels, and all signatures were manually inspected in original data by histogram visualization to confirm their validity. Identified populations were also compared with manually gated canonical T-cell subsets, to validate MEM-derived T-cell subsets (**Supplementary Fig. 6**). For rigor of final analyses, T-REX populations were filtered to exclude those comprised of 3 or fewer individuals and/or ≥85% derived from a single experimental batch, to eliminate outlier subjects and batch effects. Root-mean-square deviation (RMSD) analysis of MEM signatures was used to identify similar populations (cutoff: 90), including comparisons within one analysis (“metacluster”) and across multiple T-REX analyses. T-REX populations were converted into gates by α-convex hull and applied to all samples, including those not used in T-REX analysis, to obtain population frequencies (“alphahull”, “flowCore”, and “flowUtils” packages; F-measure >90 to ensure gate accuracy).^94–96^ Population frequencies in batch controls were compared across batches as a final quality control validation. T-REX analyses within healthy subjects (randomly split, iterated 5 times), and between healthy subjects and obese and vaccinated controls provide a reference for interpretation of COVID-19 studies (**Supplementary Figs. 4 & 5**).

#### Data Pre-Processing for BAL Analysis

Matching PBMC and BAL samples were pre-gated for live CD3+ T cells prior batch normalization using CytoNorm as previously described. Three batch control samples were used in each experiment as internal controls; one was used to train batch correction, and the others to test the validity of the correction. After batch correction, MFI peaks of each fluorophore were inspected for alignment across batches.

#### Phenograph Analysis

Normalized T-cell data were subsampled (960 events per sample) before opt-SNE dimensionality reduction and Phenograph analysis (KNN=20) using OMIQ (Dotmatics, Boston, MA, USA). Phenograph clusters with significant batch control frequency differences across experiments were excluded from the analysis for potential batch-effect errors. MEM was used to define phenotypic signatures for each cluster, and RMSD was used to identify populations that were similar to those defined by T-REX in PBMCs for each pulmonary phenotype, as previously described. Since we sought to link cells in the periphery with the lungs, Phenograph clusters that contained zero events in either all PBMC or all BAL specimens were filtered from downstream analysis. A total of 24 clusters were enriched in BAL versus PBMCs in at least 50% of subjects. An overlapping T-cell panel, including tissue-resident markers CD69 and CD103, was also employed to confirm the expression of tissue-resident markers on candidate T-cell signatures. Cells were manually gated based on MEM features and subsequently analyzed for CD103 and CD69 expression (OMIQ, Dotmatics, Boston, MA, USA).

### Univariate Statistical Analysis

Data analysis was performed in R (R, v4.2.2; RStudio, v2022.05.20-492). For analysis of cell frequencies, plasma mediators, and other continuous variables, between-group comparisons were performed using the Kruskal-Wallis test for non-normal data with Dunn’s post-test and Holm adjustment for multiple comparisons. Correlation analyses were performed using the spearman method, with Benjamini-Hochberg correction where indicated (“rstatix” package, version 0.7.2).^97^ p ≤0.05 was considered statistically significant.

### Multivariate Statistical Analysis Pipeline

Multivariate classification models were built to identify key immunological features that most effectively resolved pulmonary phenotypes from healthy. The data included in these models were plasma cytokine and autoantibody levels, and frequencies of manually gated immune subsets (spectral flow cytometry panel 2). Frequencies of T-cell populations identified with T-REX were omitted from analysis to prevent bias, as they were defined for being enriched in either post-COVID patient groups or healthy controls. AIM assay data were omitted from analysis due to low sample size. Raw values were natural log-transformed after adding 0.01 to approximately conform to normal distributions suitable for all downstream linear multivariate analyses and the generation of all correlation networks.^98^ Supervised multivariate statistical models were built using a previously described analysis pipeline.^99^ MATLAB computing environment (version R2022a, Mathworks, Natick, MA, USA) was used for multivariate analysis, and codes are available on GitHub (https://github.com/Dolatshahi-Lab/PLSR-DA; https://github.com/Dolatshahi-Lab/Elastic-Net-for-PLSR-DA).

#### Feature Selection

A minimal set of measurements was selected using elastic net regularization, in order to down-select features and prevent overfitting ^36,37^ Elastic net was performed on each dataset individually to enable inclusion of as many samples as possible. Hyperparameters were tuned to yield <20 features per data type and minimize the standard error (α, determines the relative weight of ridge and LASSO regressions in the model; stability parameter, controls how many replicates out of 100 a feature must be selected for inclusion in the final set).

#### Orthogonalized Partial Least Squares-Discriminant Analysis (OPLS-DA)

Elastic net-selected features were combined for use in OPLS-DA^38^ to identify latent variables driving discrimination of groups. To assess the goodness-of-fit, cross validation (CV) accuracy was determined using random 5-fold partitioning. To assess model robustness, empirical p-values were determined by shuffling data labels and comparing the model accuracy with 500 randomly shuffled replicates.

#### Co-correlate Networks

Given that these models select a minimal set of features that are most distinct between groups^100^, we next identified co-correlates of model-selected features that may contribute equally to group separation.^101,102^ Pearson correlation networks were generated using the same data as elastic net and OPLS-DA analyses (patient group and healthy control together)(R, v4.3.1; RStudio, v2023.09.1-494). Highly correlated features (Pearson R >0.8, FDR-adjusted p <0.05) were included in correlation networks centered around the elastic net-selected features, and visualized using Cytoscape (version 3.10.0).

#### Pulmonary Phenotype Correlation Networks

Pearson correlation networks were generated within individual pulmonary phenotypes by correlating model-selected features and co-correlates with all immune features (R, v4.3.1; RStudio, v2023.09.1-494). In addition to data used in multivariate modeling (plasma cytokines, autoantibodies, manually gated immune subsets), networks were built using T-REX population frequencies, AIM assay data (cell frequencies and secreted cytokines), and serum antibody data (SARS-CoV-2-specific IgG and total IgE) that were natural log-transformed as previously described. Features with >70% zero values were removed prior to correlation analysis. Only correlations with adjusted p-value ≤0.05 were included in network visualizations (Cytoscape, version 3.10.0). For simplicity, network visualizations in this publication focus on correlations with T-REX populations or CXCL13. Edges emanating from these nodes were manually reviewed, and correlations driven by influential points were not included in network visualizations. Complete correlation data are provided in **Supplementary Table 3**.

## Supporting information

Supplementary Figures and Table of Contents

## Acknowledgments

We would like to thank Michael Solga, MS, Director of the UVA Flow Cytometry Core Facility for spectral flow cytometry expertise. Data for this publication were generated in the UVA FCCF (RRid: SCR_017829), which is partially supported by an NCI Grant (P30-CA044579). We also thank Patcharin Pramoonjago, PhD, Director of the UVA Biorepository and Tissue Research Facility and Fabrizio Drago for coordinating the acquisition of specimens from hospitalized COVID-19 patients provided through the UVA COVID-19 Biorepository. We also thank Chintan Ramani, MD, for his role in establishing the post-clinic at UVA and facilitating enrollment into the UVA COVID-19 recovery cohort; Kyle Enfield, MD for facilitating enrollment into the COVID-19 recovery cohort; and all the clinical research support staff who performed enrollment of COVID-19 patients at UVA in both the inpatient and outpatient settings. We thank William Petri, MD PhD, for his oversight of Protocol #200110; Nathan Richards, MD and Samuel Ailsworth for assistance with antibody assays and blood draws; and Amr Atya, MD for assistance with blood draws. Finally, we would also like to thank all the participants who donated samples

## Funding

National Institutes of Health grant U01 AI100799 (SMB, GC, JMI, LMM, JAW)

National Institutes of Health grant R21 AI138077 (CB, GC, AK, LMM, JAW)

National Institutes of Health grant R56 AI178669 (CB, GC, AK, LMM, JAW)

National Institutes of Health grant T32 AI007496 (NB)

National Institutes of Health grant T32 GM145443 (SL)

Vanderbilt-Ingram Cancer Center P30 CA68485 (JMI)

University of Virginia School of Medicine GAP award (GC, LMM, JAW)

University of Virginia Global Infectious Diseases Institute award (CB, JAW)

University of Virginia Manning COVID-19 Research Fund (JW)

Henry Gustav Floren Trust, the Stanford Department of Medicine Team Science Program, and the Stanford Medicine Office of the Dean (PJU)

## Author contributions

Conceptualization: GC, SD, JMI, LMM, JAW

Methodology: GC, SD, JMI, SL, SML, LMM, PJU, JAW

Software: CEC, GC, SML, JMI, LMM

Validation: GC, SD, JMI, SML, LMM

Formal analysis, GC, SD, JMI, SL, SML, LMM, PJU, XY

Investigation: all authors

Resources: AK, JMI, CAM, JS, JMS, CAM, PJU, JAW, JMW

Data Curation: CB, GC, AK, SML, DM, LMM, JMS, JS, XY

Writing – original draft: AK, GC, SD, SL, JMI, LMM, JAW

Writing – review & editing: all authors

Visualization: CB, GC, SL, LMM, XY

Supervision: JMI, SD, JS, PJU, JAW

Project administration: JMI, JAW

Funding acquisition: CB, JMI, JS, SD, JAW

## Competing interests

JAW receives support for research unrelated to this project from Regeneron. All other authors declare that they have no competing interests.

## Data and materials availability

All data associated with this study are present in the paper, Extended Data and Supplementary Materials. All flow cytometry data and relevant metadata will be made available in FlowRepository. Materials may be shared with outside investigators subject to availability and following University of Virginia guidelines and policies.

## EXTENDED DATA

### Figures

**Extended Data Figure 1.**
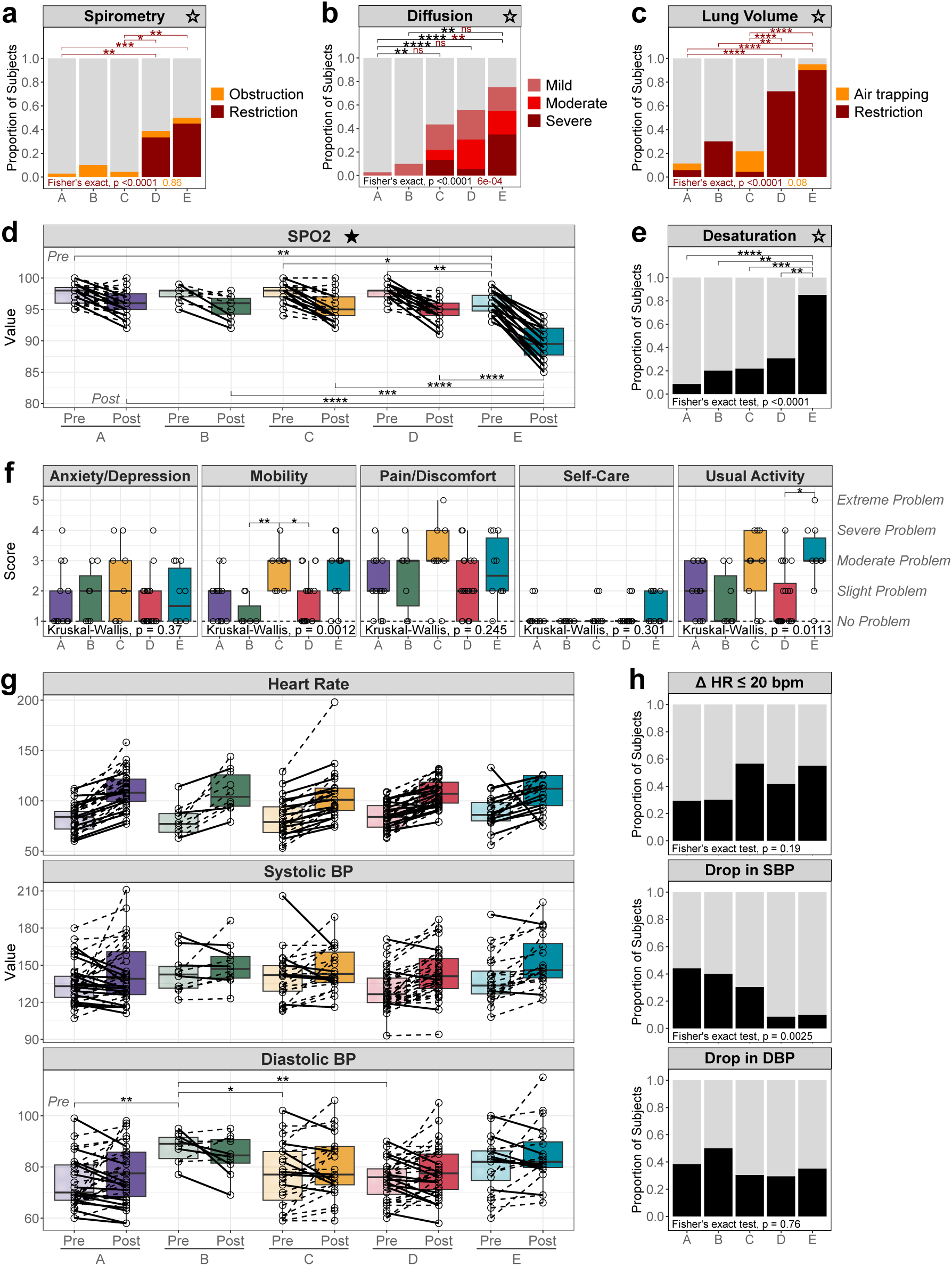
Clinical Interpretations and Additional Cardiopulmonary Measures by Patient Group. **(a–c)** Blinded interpretation of **(a)** spirometry, **(b)** diffusion capacity of the lungs for carbon monoxide (DLCO), and **(c)** lung volume assessments (124 visits). For diffusion, black statistics represent any reduction and red statistics represent severe reduction in DLCO. Cutoffs: mild ≥65%; moderate ≥50%; severe <50%. **(d–e)** Blood oxygen saturation during 6MWT (124 visits), including **(d)** change after test (solid lines denote desaturation), and **(e)** the proportion of subjects who desaturated (≥ 3% decrease). **(f)** Patient reported quality of life scores, where available. (EQ-5D-5L, 59 visits). **(g–h)** Heart rate (HR) and blood pressure (BP) responses during 6MWT (≥ 121 visits), including **(g)** change after test (solid lines denote abnormal responses), and **(h)** proportion of abnormal responses by patient group. HR, change ≤ 20 bpm; Systolic and diastolic BP, any decrease in mmHg. ★, measures used to derive clinical phenotypes; ☆, categorizations also derived from ★ measures. Fisher’s exact test (***a–c, e, h***), and Kruskal-Wallis with Dunn’s post-hoc test and Holm adjustment (***d & g:*** *Pre & Post* analyzed separately; ***f***). \**P* ≤ 0.05; \*\**P* ≤ 0.01; \*\*\**P* ≤ 0.001; \*\*\*\**P* ≤ 0.0001.

**Extended Data Figure 2.**
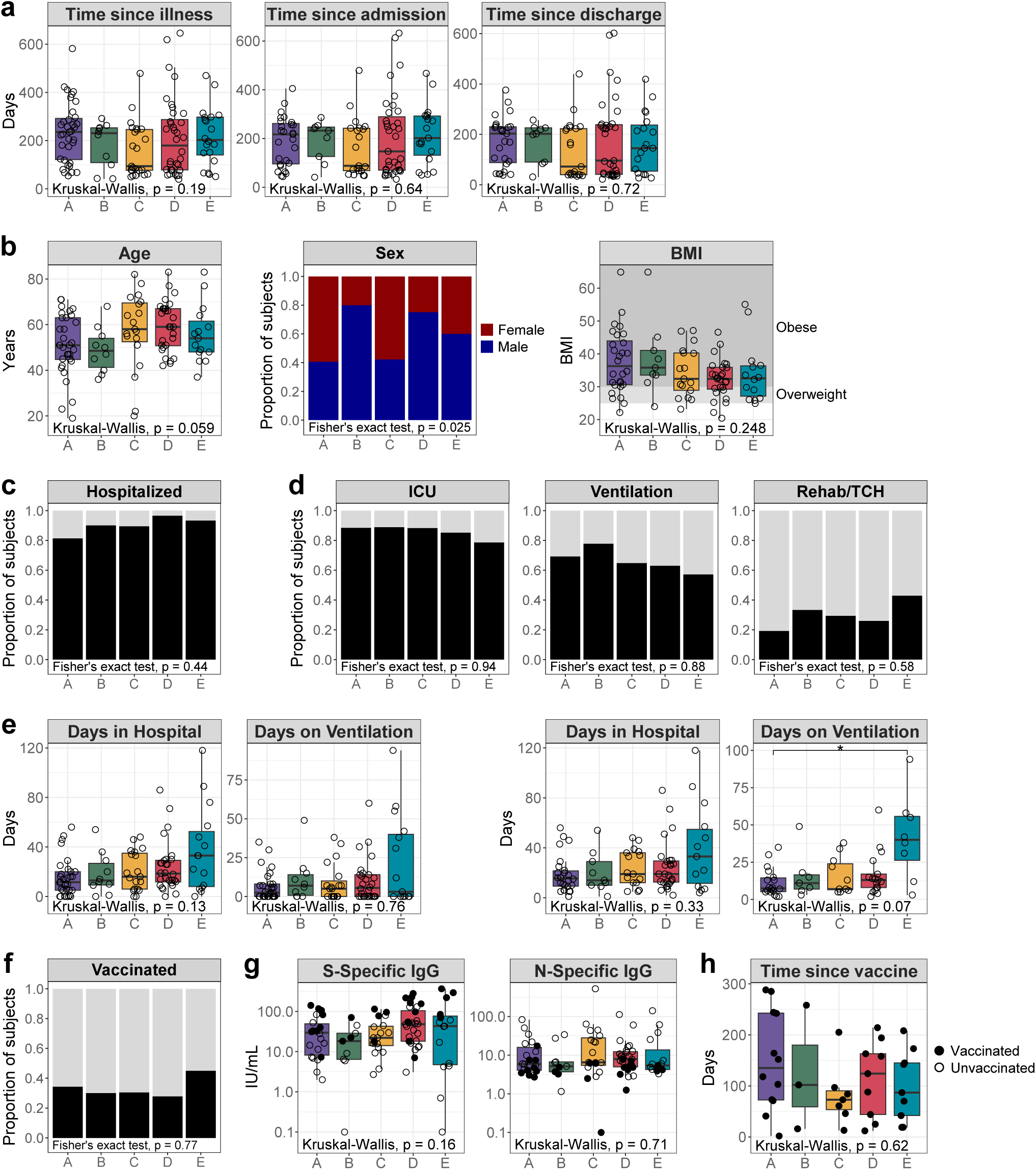
Relevant Biological, Clinical, and Vaccination Variables by Patient Group. **(a)** Time since initial illness (124 visits), and hospital admission and discharge (hospitalized only, 111 visits). **(b)** Age and sex (104 patients) and BMI (95 patients) at the time of initial illness. **(c)** Proportion of patients requiring hospitalization (*n* = 104). **(d)** Proportion of hospitalized patients requiring ICU care, mechanical ventilation, or discharge to a rehabilitation facility/transitional care hospital (TCH) (*n* = 93). **(e)** Duration of hospital stay and mechanical ventilation for all patients (*left,* 104 patients), and for those who were hospitalized or ventilated, respectively (*right*, 93 and 61 patients, respectively). **(f)** Proportion of patients that were vaccinated at the time of follow-up visit (124 visits). **(g)** SARS-CoV-2 original strain spike (S)- and nucleocapsid (N)-specific IgG (124 visits). **(h)** Days elapsed since vaccination according to patient cluster (40 visits). Vaccinated patients denoted by filled symbols (***g, h***). Kruskal-Wallis with Dunn’s post-hoc test and Holm adjustment (continuous variables), and Fisher’s exact test (categorical variables). Duplicate patients within the same cluster were removed from analysis (***b, c, d, e***). \**P* ≤ 0.05.

**Extended Data Figure 3.**
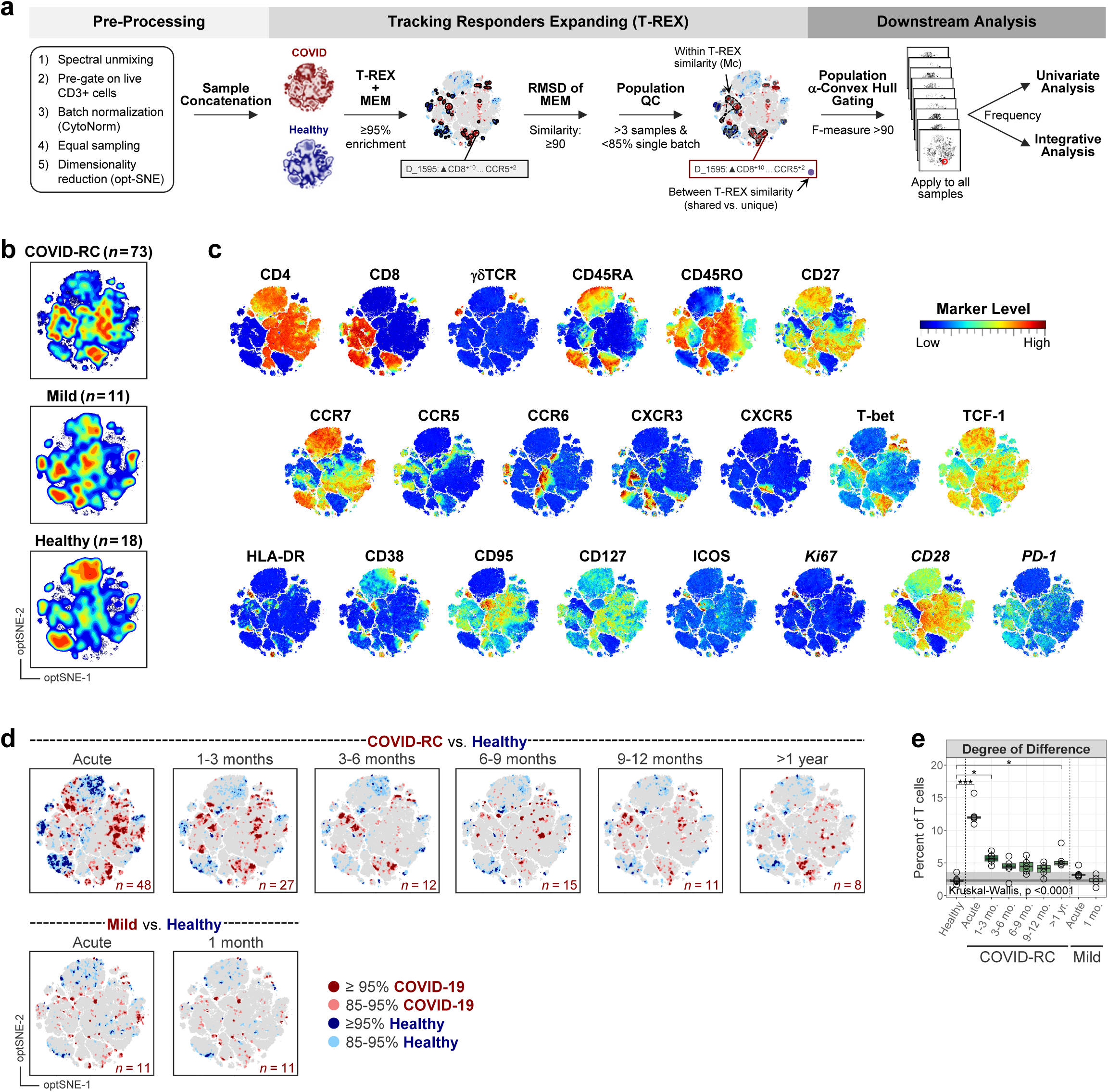
T-Cell Marker Expression and Longitudinal Monitoring After COVID-19 Illness. **(a)** Data analysis pipeline for Tracking Responders Expanding (T-REX) analysis of T cells. Root-mean-square deviation (RMSD) of T-cell signatures is used to: (i) group similar populations within a single T-REX analysis (metacluster, Mc; denoted on plots by dashed outlines); and (ii) compare populations between multiple T-REX analyses (ie. comparing across pulmonary phenotypes A-E; denoted in population labels by colored dots). **(b)** opt-SNE dimensionality reduction of CD3+ T cells following COVID-19 illness (COVID-RC and mild COVID-19), and in healthy groups. **(c)** Marker expression plots. Markers denoted in italics were not utilized in dimensionality reduction. **(d)** T-REX plots for COVID-RC and mild COVID-19 groups over time (time since admission [hospitalized] or start of illness [outpatient]), as compared with healthy controls (*n* = 18). Red populations are increased in COVID; blue populations are increased in healthy (decreased in COVID). **(e)** T-REX Degree of Difference in COVID-19 versus healthy groups over time (percent of cells in regions of 95% enrichment). Analyses were iterated excluding single batches (*n* = 5). Shading shows the range (*light*) and IQR (*dark*) of healthy subjects (see **Supplementary Fig. 4**). Kruskal-Wallis with Dunn’s post-hoc test and Holm adjustment, comparison with healthy only. \**P* ≤ 0.05; \*\**P* ≤ 0.01; \*\*\**P* ≤ 0.001.

**Extended Data Figure 4.**
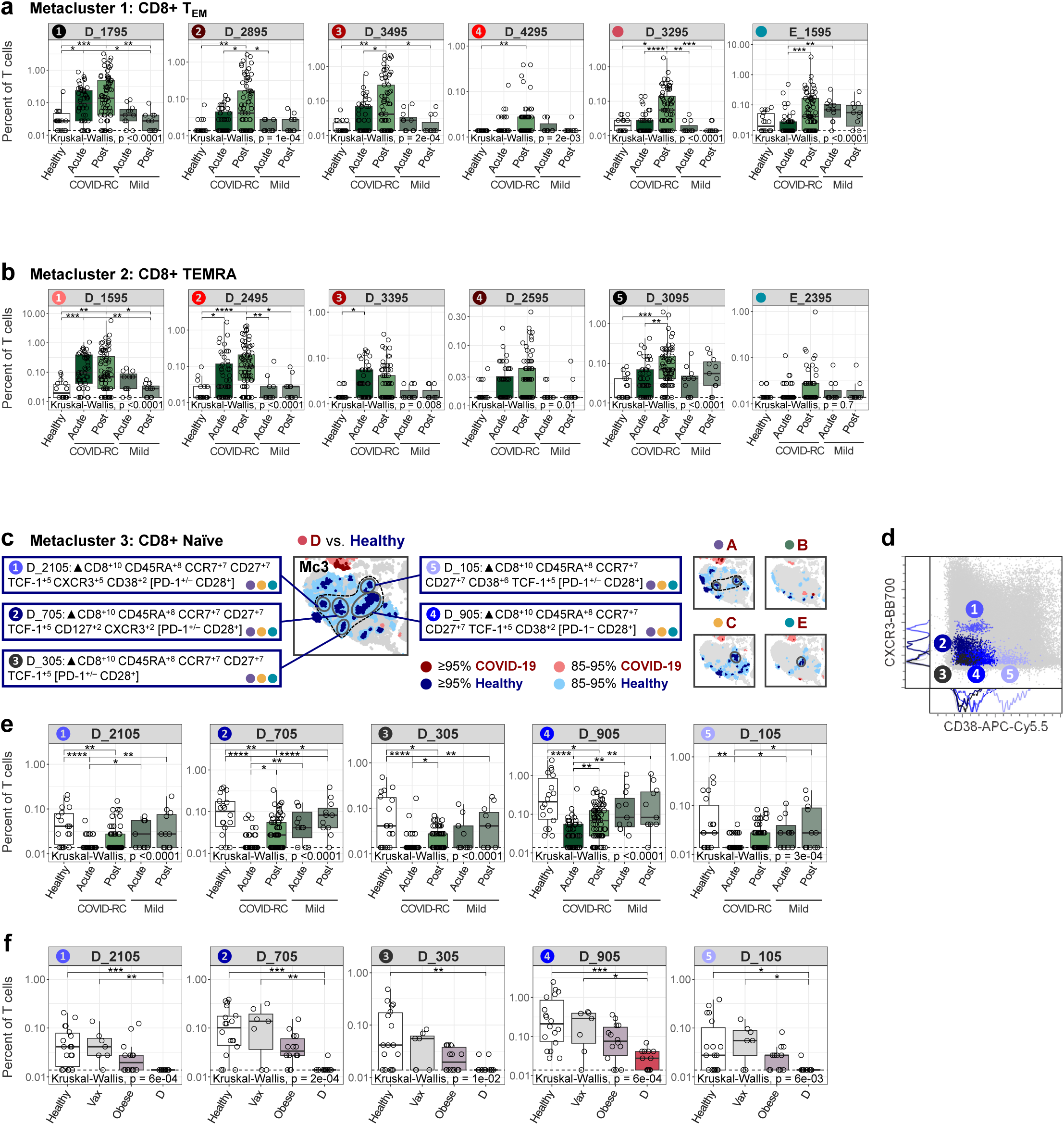
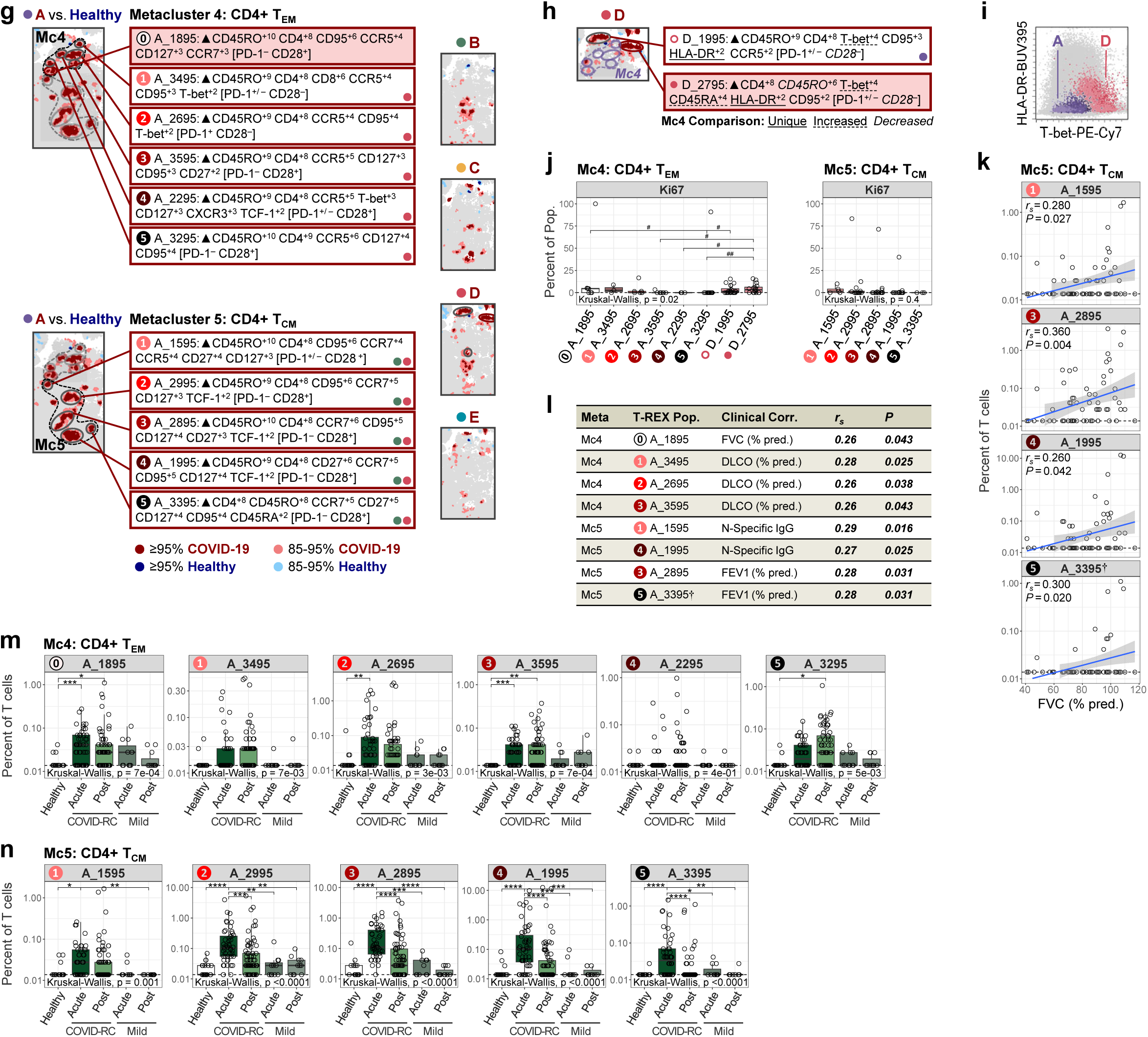
Dysregulation of Mc1, Mc2, and Mc3 in Restrictive Lung Disease. **(a & b)** Frequencies of **(a)** Mc1 and **(b)** Mc2 and corresponding unique D and E populations across acute and post-COVID time points in COVID-RC and mild COVID groups (*n* = 161; ≥11 per group). **(c)** CD8+ naïve T-cell populations in Mc3 with corresponding MEM labels, as identified by T-REX of group D (*n* = 12) vs. healthy controls (*n* = 18) and RMSD (similarity cutoff, 90). Colored dots in MEM label boxes denote signature similarity across patient groups. PD-1 and CD28 (measured in 4 of 5 batches) were excluded from T-REX analyses and MEM labels, and expression is summarized in brackets. **(d)** Expression of CXCR3 and CD38 across Mc3 populations. Concatenated data, total T cells shown in gray. **(e-f)** Frequencies of Mc3 populations across **(e)** acute and post-COVID time points in COVID-RC and mild COVID groups (*n* = 161; ≥11 per group), and **(f)** healthy, vaccinated, and obese control groups (*n* = 52; ≥7 per group). Kruskal-Wallis with Dunn’s post-hoc test and Holm adjustment (***a, b, e, f***). \**P* ≤ 0.05; \*\**P* ≤ 0.01; \*\*\**P* ≤ 0.001 ; \*\*\*\**P* ≤ 0.0001. **Mc4 and Mc5 Link to Improved Lung Function. (g)** CD4+ T_EM_ and T_CM_ populations in Mc4 and Mc5, respectively, as identified by T-REX of group A (*n* = 12) vs. healthy controls (*n* = 18) and RMSD. A signature unique to A is shown in a shaded box. **(h)** Group D CD4+ T_EM_ populations that are adjacent to Mc4, including shared (with group A) and unique (shaded box) signatures. MEM distinguishing features are highlighted. **(i)** Scatter plot comparison of T-bet and HLA-DR expression in group A (Mc4) and group D T_EM_ populations (concatenated data, total T cells depicted in gray). **(j)** Cell proliferation (Ki67) across Mc4 and related D populations (*left; n* = 42; ≥3 per pop) and Mc5 (*right; n* = 51; ≥3 per pop). **(k)** Significant spearman correlations between Mc5 frequencies and FVC in the COVID-COVIDRC (*n* = 61). **(l)** Significant spearman correlations between Mc4 and Mc5 frequencies and other measures in the COVID-RC (*n* ≥ 61). **(m & n)** Frequencies of **(m)** Mc4 and **(n)** Mc5 populations across acute and post-COVID time points in COVID-RC and mild COVID groups (*n* = 161; .11 per group). PD-1 and CD28 (measured in 4 of 5 batches) were excluded from T-REX analyses and MEM labels, and expression is summarized in brackets **(g, h)**. Colored dots in MEM label boxes denote signature similarity across patient groups (similarity cutoff, 90)*(g, h)*. Kruskal-Wallis with Dunn’s post-hoc test and Holm adjustment ***(j, m, n)***. *P ≤ 0.05; **P ≤ 0.01; ***P ≤ 0.001 ; ****P ≤ 0.0001 (^#^, unadjusted).

**Extended Data Figure 5.**
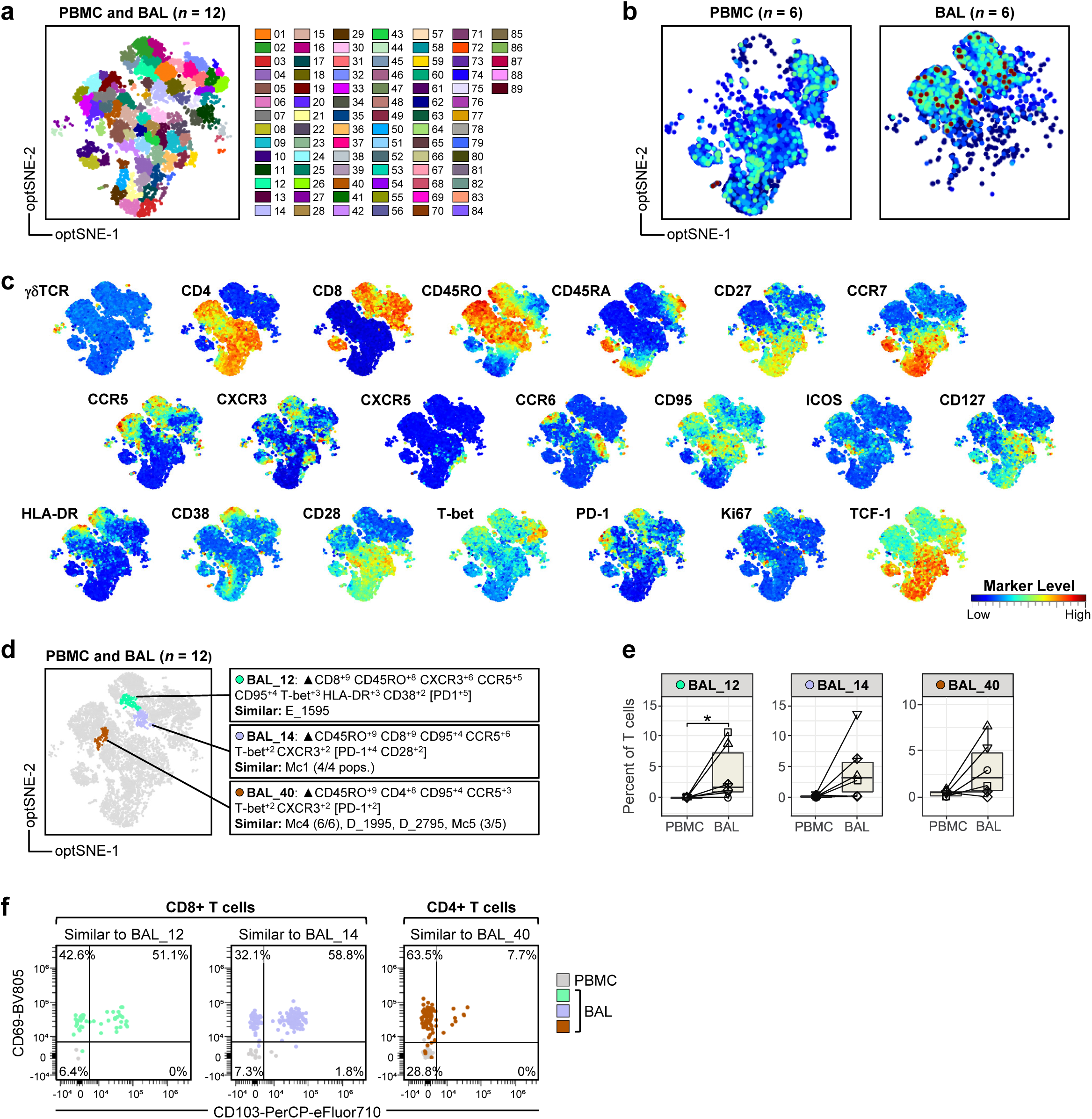
Paired Analysis of T Cells in the Blood and Airways of Patients with Prior COVID-19. **(a-c)** opt-SNE dimensionality reduction of CD3+ T cells in paired PBMC and BAL samples from patients with a history of COVID-19 (*n* = 6). **(a)** Phenograph analysis. **(b)** Cell density plots. **(c)** Marker expression plots. **(d**) MEM signatures of phenograph populations that are enriched in BAL and are similar to E_1595 (BAL_12), Mc1 (BAL_14), or Mc4 (BAL_40) T-REX populations (pops), as determined by RMSD analysis (cutoff, 90). **(e)** Frequency of phenograph populations in paired PBMC and BAL samples from patients with prior COVID-19 (*n* = 6). **(f)** Representative plots confirming CD69 and CD103 expression on CD8+ and CD4+ T-cell populations similar to those highlighted in ***d*** and ***e***, gated according to MEM signatures.

**Extended Data Figure 6.**
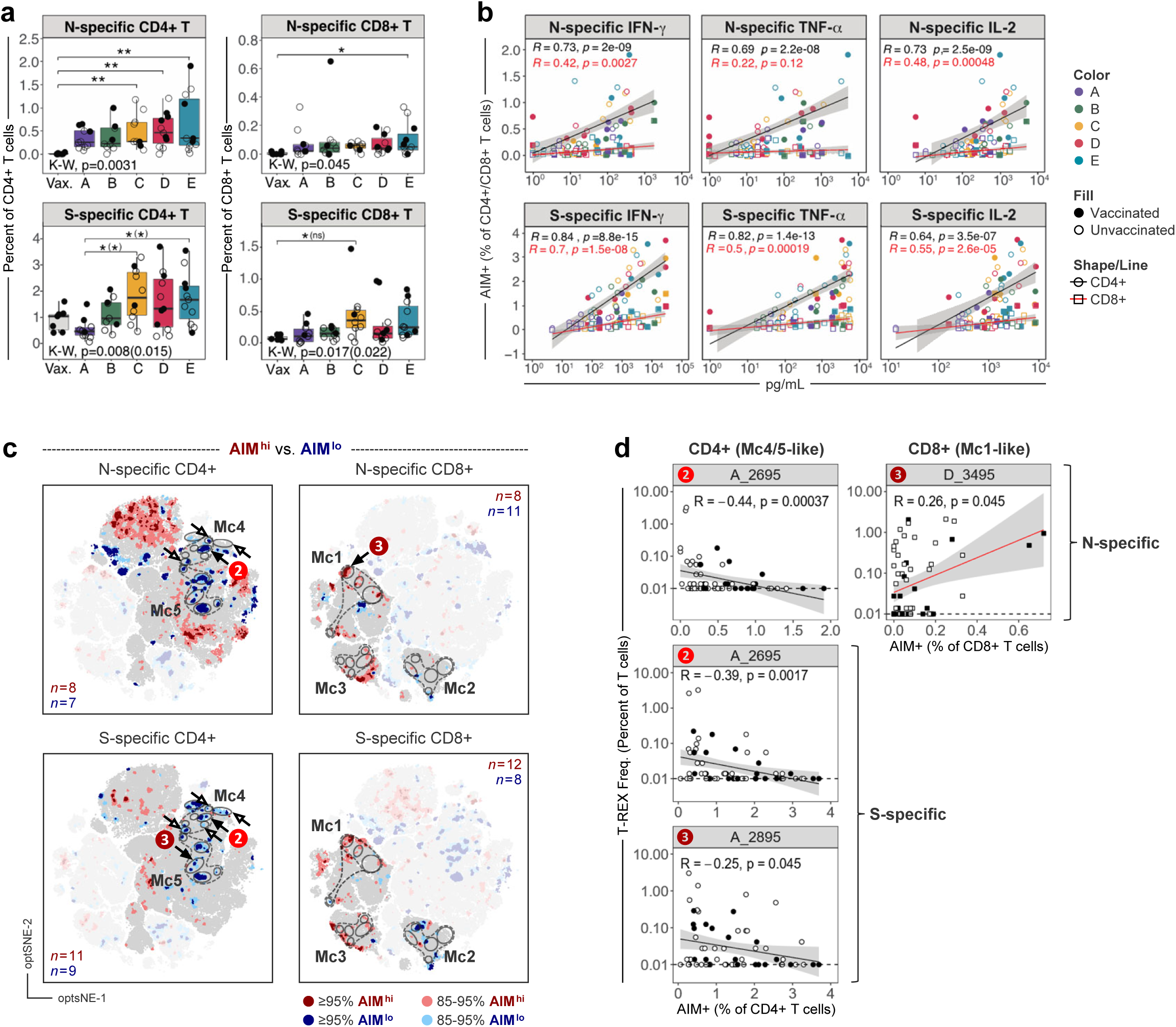
SARS-CoV-2-specific T Cells are Elevated in Patients with Persistent Respiratory Complications After COVID-19. **(a)** Frequencies of Nucleocapsid (N) and Spike (S)-specific CD4+ (OX40+CD137+) and CD8+ (CD69+CD137+) T cells detected by AIM assay (*n* ≥ 57, ≥7 per group). Filled circles indicate samples obtained after SARS-CoV-2 vaccination. Kruskal-Wallis (K-W) with Dunn’s post-hoc test and Holm adjustment. Statistical comparisons excluding vaccinated COVID-RC subjects are given in parentheses (*n* = 41, ≥6 per group). **(b)** Spearman correlation of N- and S-specific CD4+ (black line, circles) and CD8+ (red line, squares) T cells and cytokines in corresponding culture supernatants (*n* = 51 each). **(c)** T-REX comparison of subjects with the highest and lowest AIM+ T-cell frequencies within the COVID-RC, regardless of pulmonary phenotype or vaccination status. Non-corresponding CD4+ or CD8+ T cells are faded. Metaclusters as defined in Figure 2 are outlined for reference. Arrows denote populations that correlate with AIM frequencies (filled, correlations shown in ***d***). N-specific CD4+: hi ≥1%, lo ≤0.1%; N-specific CD8+: ≥0.2%, ≤0.01%; S-specific CD4+: ≥2.5%, ≤0.4%; S-specific CD8+: ≥0.5%, ≤0.05%. **(d)** Spearman correlations of AIM+ T-cell frequencies with corresponding T-REX population frequencies (*n* ≥ 60). Vax., vaccinated healthy subjects (6 months post-vaccination). For all AIM assays, values are background subtracted using unstimulated conditions (lower limit zero). \**P* ≤ 0.05; \*\**P* ≤ 0.01; \*\*\*\**P* ≤ 0.0001.

**Extended Data Figure 7.**
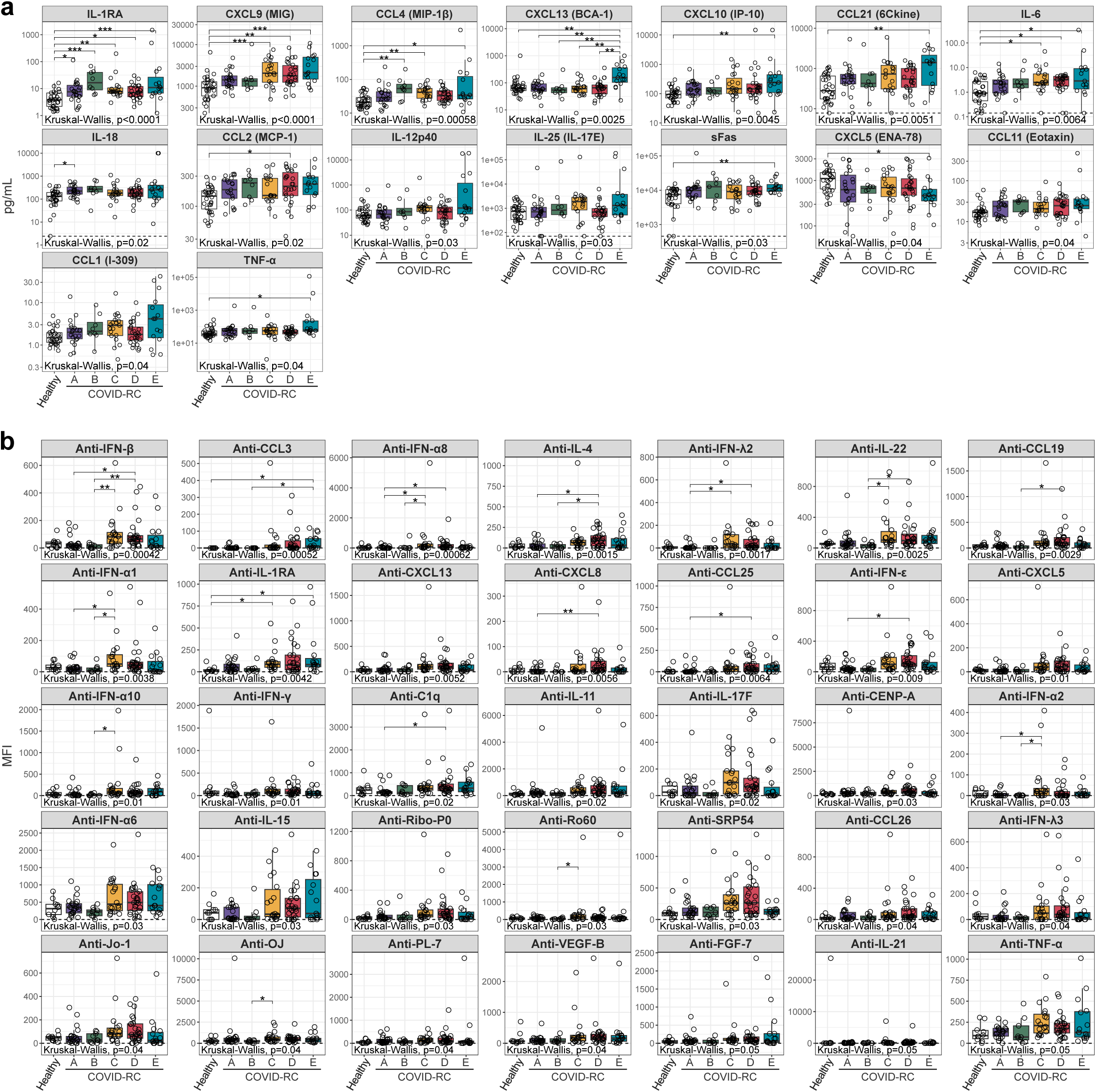
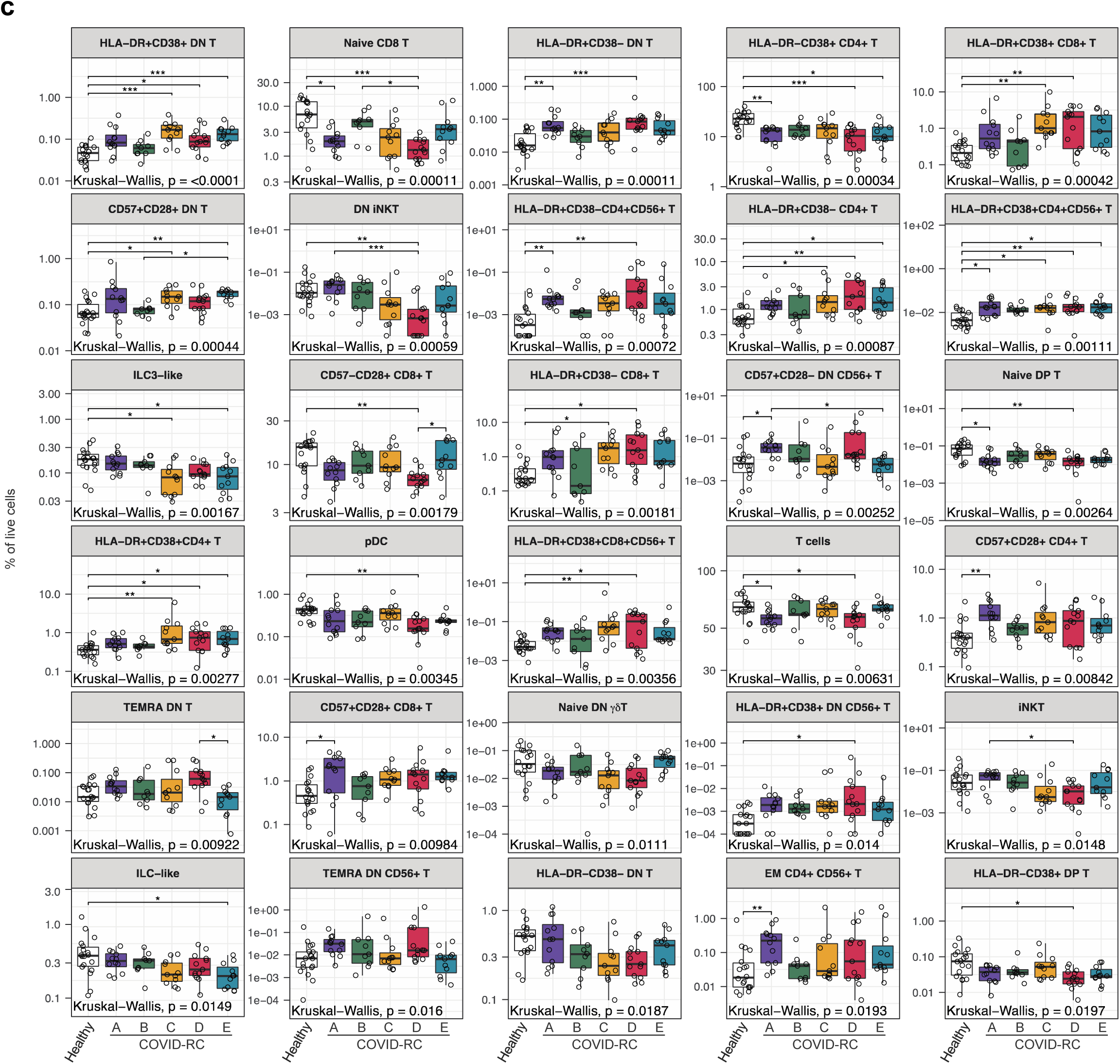
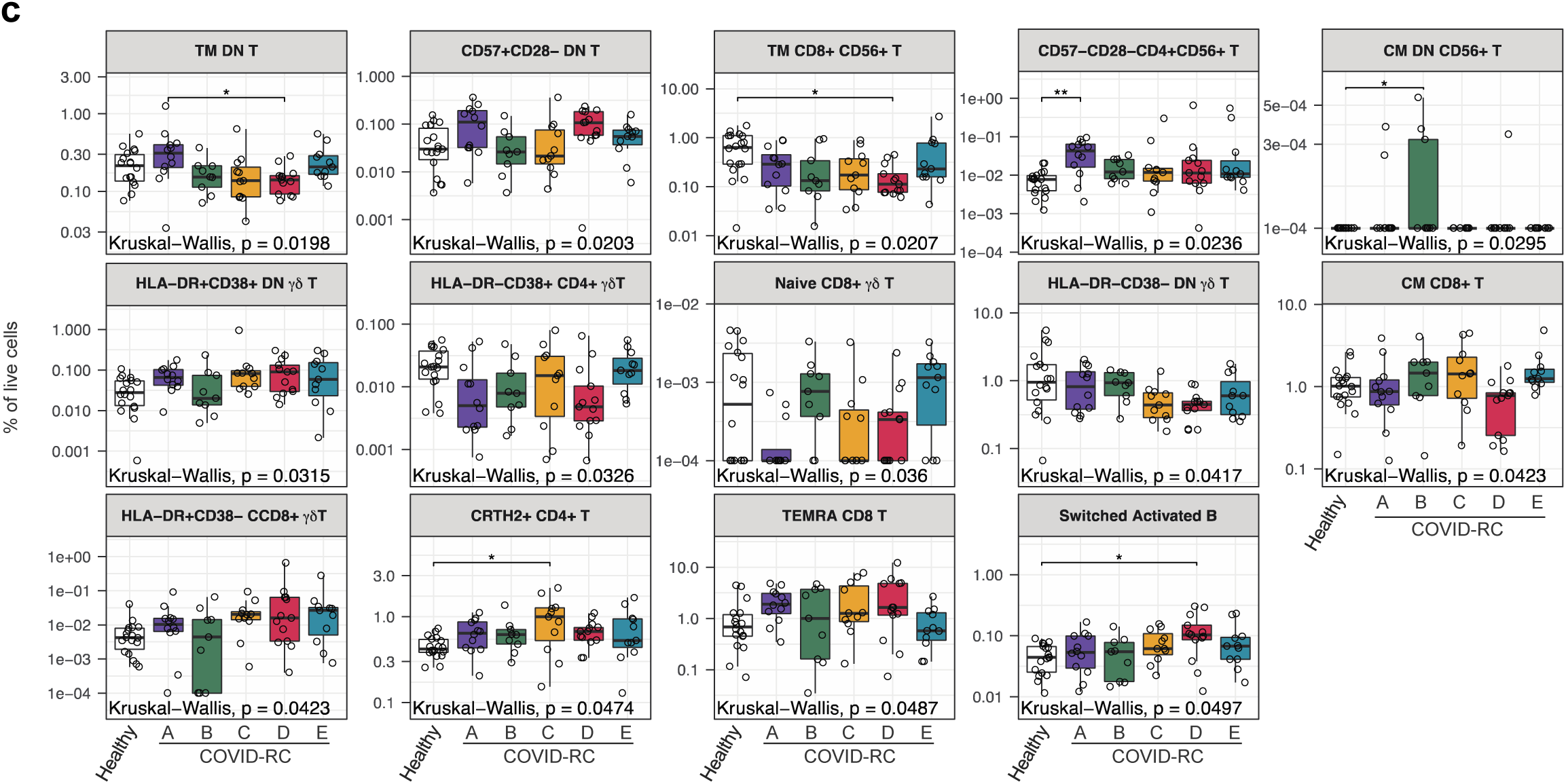
Broad Monitoring of Cytokines, Auto-antibodies, and Immune Cells Reveals Distinct Profiles Between Patient Groups. Levels of **(a)** plasma cytokines (pg/mL; *n* ≥ 120, ≥ 8 per group), **(b)** auto-antibodies (MFI; *n* = 100, ≥ 9 per group), and **(c)** manually gated canonical immune cell types and subsets (percent of live cells; *n* = 74, ≥ 9 per group). Kruskal-Wallis (K-W) with Dunn’s post-hoc test and Holm adjustment. Only measures with K-W *P* ≤ 0.05 are shown. \**P* ≤ 0.05; \*\**P* ≤ 0.01; \*\*\**P* ≤ 0.001; \*\*\*\**P* ≤ 0.0001.

**Extended Data Figure 8.**
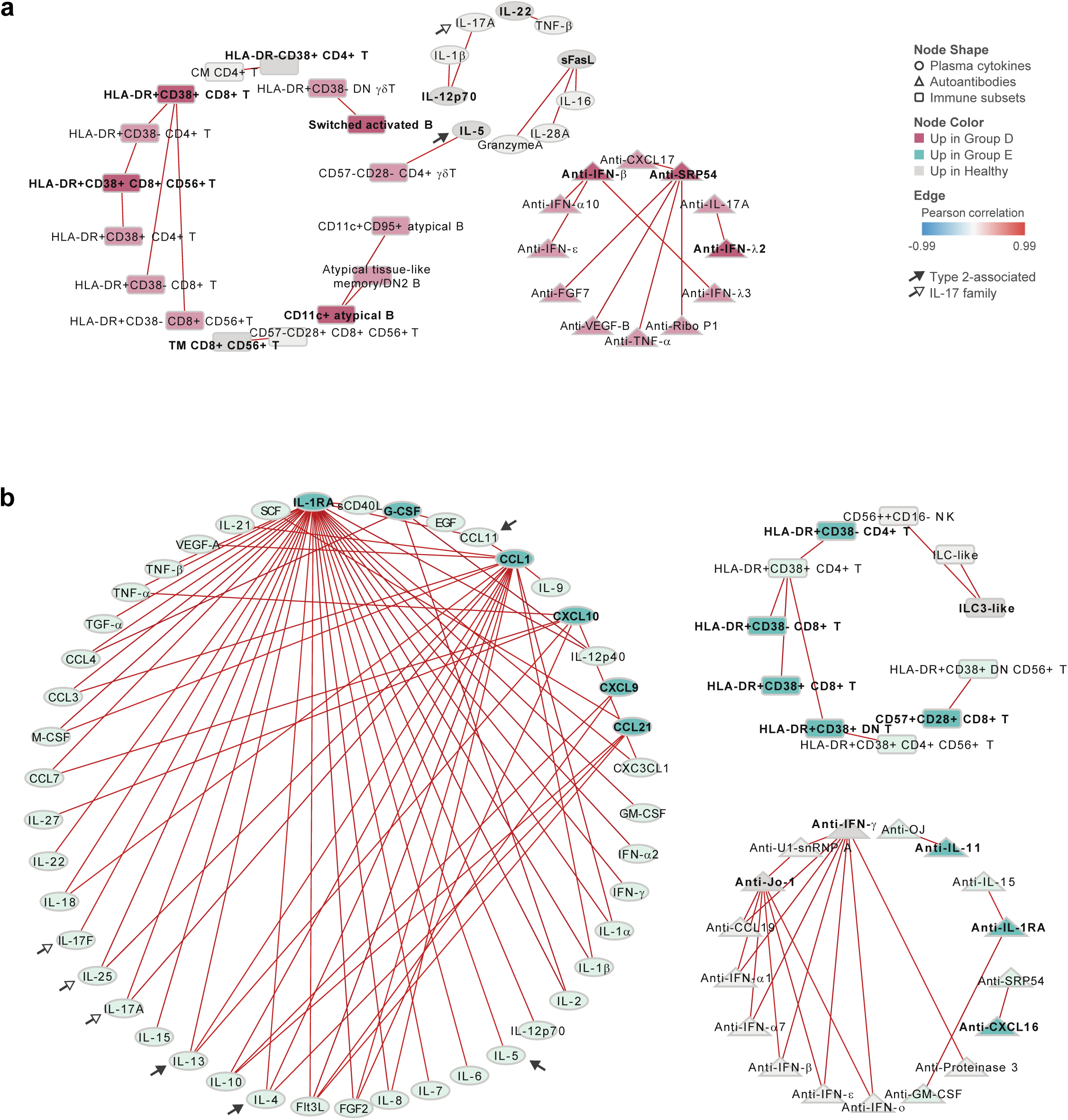
Co-correlate Networks of Restrictive Lung Disease Phenotypes D and E. **(a & b)** Pearson correlation networks depicting features that are highly correlated (|*R*| >0.8) with elastic net-selected features. Correlations were run in **(a)** group D (*n* = 12), or **(b)** group E (*n* = 11), together with healthy controls (*n* = 10). Node shape denotes data type, and color denotes elastic net-selected features (dark) and co-correlates (light); edge color represents Pearson *R*. Type 2-associated and IL-17 family mediators are highlighted by arrows.

**Extended Data Figure 9.**
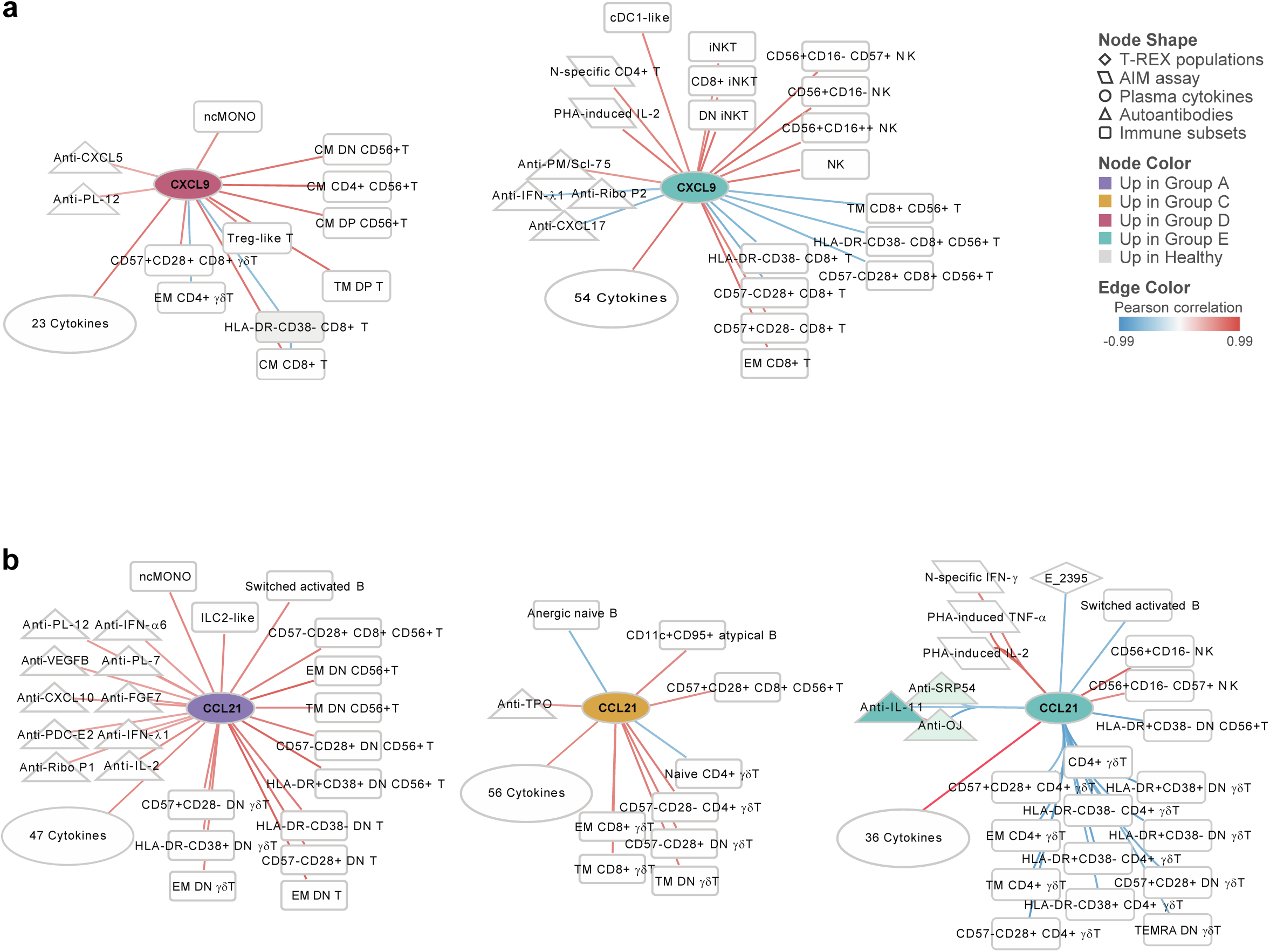
Correlation Networks of CXCL9 and CCL21 Diverge Across Pulmonary Phenotypes. Pearson correlation networks generated within patient groups (*n* ≥ 9) depicting only those features that significantly correlate with **(a)** CXCL9, or **(b)** CCL21. Data utilized in correlation analyses include T-REX population frequencies, AIM assay data (cell frequencies and secreted cytokines), and SARS-CoV-2-specific IgG, in addition to data utilized in elastic net and OPLS-DA modeling. For readability, cytokine correlations are summarized. The full list of cytokine correlations can be found in **Supplementary Table 3.** Node shape denotes data type, and color denotes elastic net-

**Extended Data Figure 10.**
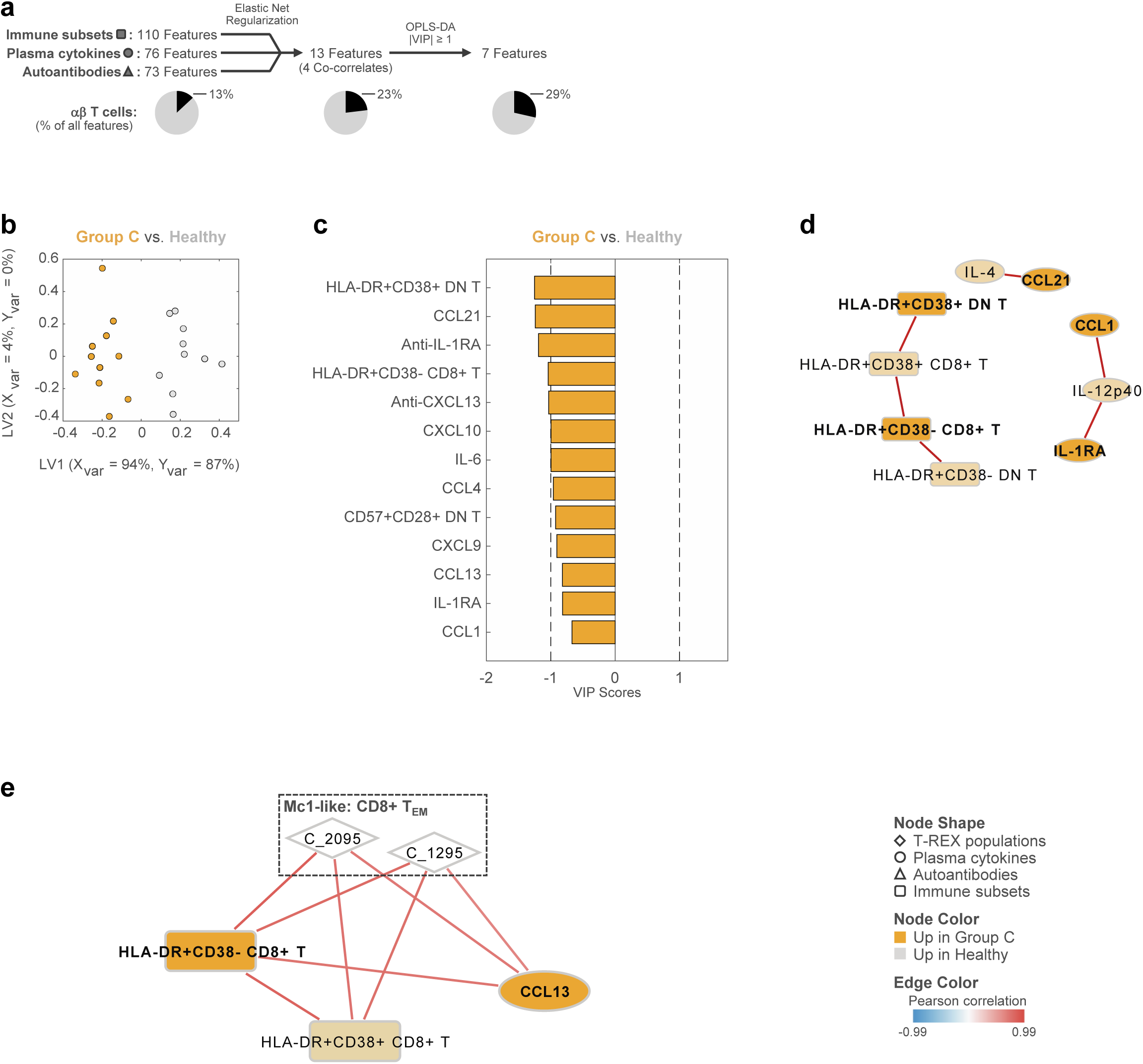
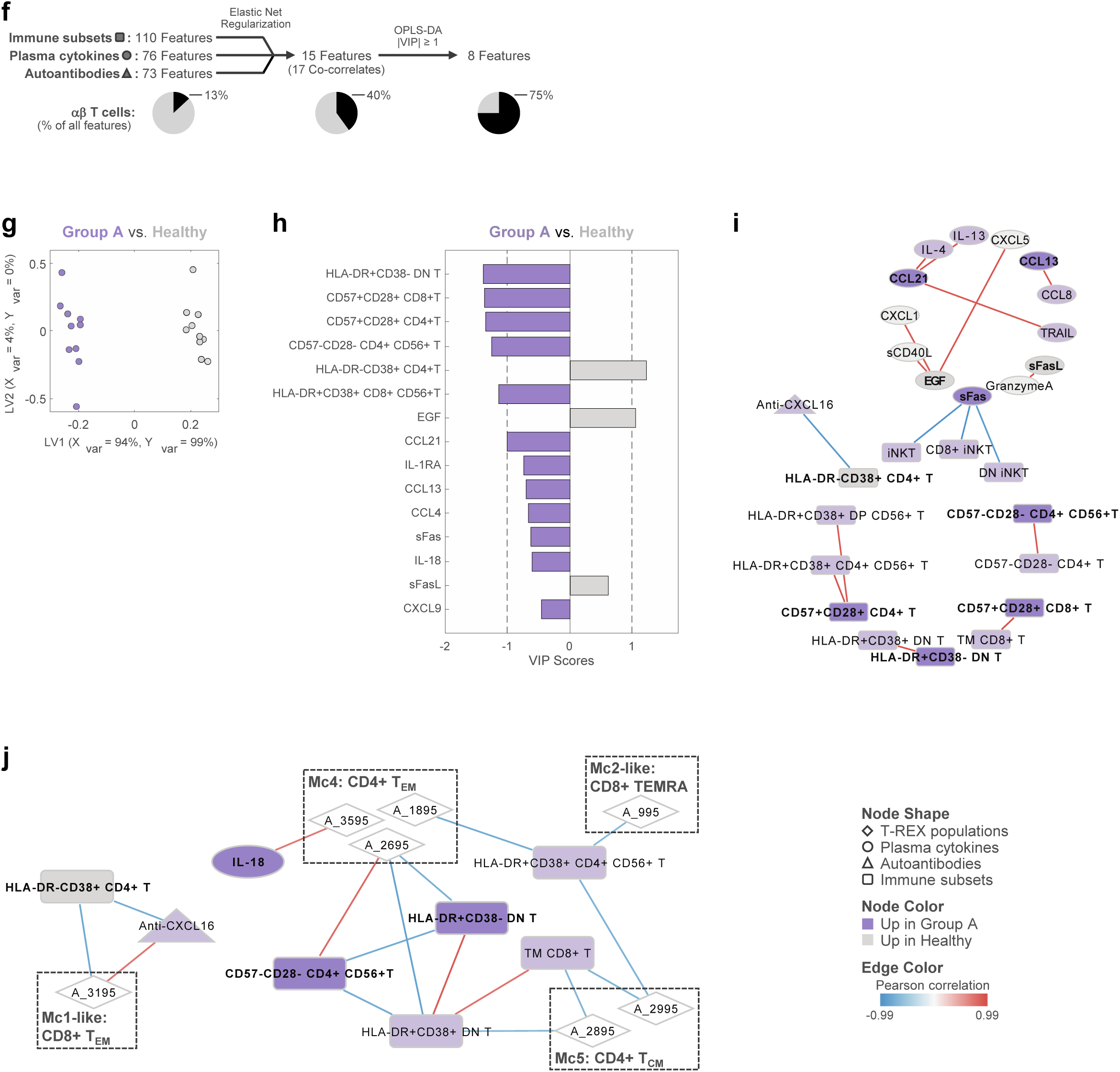
Immune Features That Discriminate Group C from Healthy Controls. **(a) I**ntegrative analysis pipeline for the generation of an OPLS-DA model that discriminates group C from healthy controls. Elastic net feature selection was run separately for each dataset before performing combined OPLS-DA. The percent of features that correspond to αβ T cells (CD3+γδTCR-) are denoted. **(b)** OPLS-DA score plot depicting the model of separation for group C (*n* = 11) versus healthy controls (*n* = 10), where each point represents one sample. Use of an orthogonalized approach ensured that latent variable 1 (LV1) captures between-group separation while simultaneously capturing 87% of the Y-variation, while LV2 captured variances that did not contribute to this separation. 5-fold cross-validation (CV) resulted in 100% CV accuracy. The model significantly outperformed models based on shuffled group labels (permutation testing, Wilcoxon *p*=0.002). **(c)** Ordered Variable Importance in Projection (VIP) scores of all elastic net-selected features for the discrimination of group C from healthy controls. **(d)** Pearson correlation network depicting features that are highly correlated (|*R*| >0.8) with elastic net-selected features. Correlations were run in group C (*n* = 11) and healthy controls (*n* = 10) together. **(e)** Pearson correlation network generated within group C (*n* ≥ 9) depicting only those features that significantly correlate with T-REX populations identified in group C. Data utilized in correlation analyses include T-REX population frequencies, AIM assay data (cell frequencies and secreted cytokines), and SARS-CoV-2-specific IgG, in addition to data utilized in elastic net and OPLS-DA modeling. Node shape denotes data type, and color denotes elastic net-selected features (dark) and co-correlates (light); edge color represents Pearson *R* (*P_adj_* ≤0.05). **(f) I**ntegrative analysis pipeline for the generation of an OPLS-DA model that discriminates group A from healthy controls. Elastic net feature selection was run separately for each dataset before performing combined OPLS-DA. The percent of features that correspond to αβ T cells (CD3+γδTCR-) are denoted. **(g)** OPLS-DA score plot depicting the model of separation for group A (*n* = 10) versus healthy controls (*n* = 10), where each point represents one sample. Use of an orthogonalized approach ensured that LV1 captures between-group separation while simultaneously capturing 99% of the Y-variation, while LV2 captured variances that did not contribute to this separation. 5-fold cross-validation (CV) resulted in 100% CV accuracy. The model significantly outperformed models based on shuffled group labels (permutation testing, Wilcoxon *p*=0.002). **(h)** Ordered Variable Importance in Projection (VIP) scores of all elastic net-selected features for the discrimination of group A from healthy controls. **(i)** Pearson correlation network depicting features that are highly correlated (|*R*| >0.8) with elastic net-selected features. Correlations were run in group A (*n* = 10) and healthy controls (*n* = 10) together. **(j)** Pearson correlation network generated within group A (*n* ≥ 10) depicting only those features that significantly correlate with T-REX populations identified in group A. Data utilized in correlation analyses include T-REX population frequencies, AIM assay data (cell frequencies and secreted cytokines), and SARS-CoV-2-specific IgG, in addition to data utilized in elastic net and OPLS-DA modeling. Node shape denotes data type, and color denotes elastic net-selected features (dark) and co-correlates (light); edge color represents Pearson *R* (*P_adj_* ≤0.05).

