## Supplementary Table 4 for "Distinct Type 1 Immune Networks Underlie the Severity of Restrictive Lung Disease after COVID-19"

**Supplementary Table 4. Spectral Flow Cytometry Panels**

| **Panel 1: T cell deep immunophenotyping ^#^** | | | | |
| --- | --- | --- | --- | --- |
| **MARKER** | **FLUOROPHORE** | **CLONE** | **CATALOG #** | **COMPANY** |
| Live/Dead | Blue | NA | L23105 | THERMO FISHER |
| CCR5 | BUV 737 | 2D7 | 565293 | BD |
| CCR6 | BUV661 | 11A9 | 750696 | BD |
| CXCR3 | BB700 | 1C6 | 566532 | BD |
| CXCR5 | BV421 | J252D4 | 356920 | BIOLEGEND |
| CCR7 | APC/Fire 750 | G043H7 | 353246 | BIOLEGEND |
| HLA-DR | BUV395 | G46-6 | 564040 | BD |
| CD4 | BUV496 | OKT4 | 750980 | BD |
| CD8 | BUV805 | RPA-T8 | 749366 | BD |
| ICOS | Pacific Blue | C398.4 | 313522 | BIOLEGEND |
| TCRγδ | BV480 | 11F2 | 746498 | BD |
| CD27 | BV570 | O323 | 302825 | BIOLEGEND |
| CD45RA | BV605 | HI100 | 304133 | BIOLEGEND |
| CD28 | BV650 | CD28.2 | 302946 | BIOLEGEND |
| CD127 | BV750 | HIL-7R-M21 | 747089 | BD |
| PD-1 | BV785 | EH12.2H7 | 329930 | BIOLEGEND |
| CD45RO | BB515 | UCHL1 | 564529 | BD |
| CD14 | FITC | M5E2 | 301804 | BIOLEGEND |
| CD19 |  | HIB19 | 302206 | BIOLEGEND |
| CD3 | Spark Blue 550 | SK7 | 344852 | BIOLEGEND |
| CD95 | BV711 | DX2 | 305644 | BIOLEGEND |
| CD38 | APC-Cy 5.5 | HIT2 | MHCD3819 | LIFE TECHNOLOGIES |
| T-bet | PECY7 | 4B10 | 644824 | BIOLEGEND |
| Ki-67 | APC | 20RAJ1 | 17-5699-42 | INVITROGEN |
| TCF-1 | AF647 | C63D9 | 6709S | CELL SIGNALING |
| **Panel 2: Main Immune Subsets** | | | | |
| **MARKER** | **FLUOROPHORE** | **CLONE** | **CATALOG #** | **COMPANY** |
| CCR7 | BV480 | 3D12 | 566099 | BD |
| Live/Dead | Blue | NA | L23105 | THERMO FISHER |
| CD21 | BUV395 | B-ly4 | 740288 | BD |
| CD15 | BUV496 | W6D3 | 741187 | BD |
| CD27 | BUV661 | O323 | 751680 | BD |
| HLA-DR | BUV805 | G46-6 | 748338 | BD |
| TCR Vα24-Jα18 | BV421 | 6B11 | 342916 | BIOLEGEND |
| CD123 | SB436 | 6H6 | 62-1239-42 | THERMO FISHER |
| CD16 | Pacific Blue | 3G8 | MHCD1628 | THERMO FISHER |
| CD14 | BV510 | M5E2 | 301842 | BIOLEGEND |
| CD8 | BV570 | RPA-T8 | 301038 | BIOLEGEND |
| CD1c | BV605 | L161 | 331538 | BIOLEGEND |
| CD56 | BV650 | HCD56 | 318344 | BIOLEGEND |
| CD19 | BV711 | SJ25C1 | 363022 | BIOLEGEND |
| CD4 | BV750 | SK3 | 344644 | BIOLEGEND |
| CD28 | BV785 | CD28.2 | 302950 | BIOLEGEND |
| CD11c | BB515 | BU-15 | 566835 | BD |
| CD45RA | Alexa 488 | HI100 | 304114 | BIOLEGEND |
| CD3 | Alexa 532 | UCHT1 | 58-0038-42 | THERMO FISHER |
| CD33 | PerCP | WM53 | A15804 | THERMO FISHER |
| CD11b | PerCP-Cy5.5 | ICRF44 | 301328 | BIOLEGEND |
| TCRγδ | BB700 | 11F2 | 745944 | BD |
| CD117 | PE | 104D2 | 313204 | BIOLEGEND |
| IgD | PE-Dazzle | IA6-2 | 348240 | BIOLEGEND |
| CD95 | PE-Cy5 | DX2 | 305610 | BIOLEGEND |
| CD25 | PEAF700 | 3G10 | MHCD2524 | THERMO FISHER |
| CRTH2 | PE-Cy7 | BM16 | 350118 | BIOLEGEND |
| TCR Vβ11 | APC | C21 | A66905 | BECKMAN COULTER |
| CD57 | eFluor 660 | TB01 | 50-0577-42 | THERMO FISHER |
| CD127 | APC-R700 | HIL-7R-M21 | 565185 | BD |
| CD38 | APCfire810 | HIT2 | 303550 | BIOLEGEND |
| **Panel 3: Activation-induced marker (AIM)** | | | | |
| **MARKER** | **FLUOROPHORE** | **CLONE** | **CATALOG #** | **COMPANY** |
| Live/dead | Blue | NA | L23105 | THERMO FISHER |
| CD3 | SB550 | SK7 | 344852 | BIOLEGEND |
| CD4 | BUV496 | OKT4 | 750980 | BD |
| CD8 | BUV805 | RPA-T8 | 749366 | BD |
| CD45RA | BV605 | HI100 | 304133 | BIOLEGEND |
| CCR7 | APC fire750 | G043H7 | 353246 | BIOLEGEND |
| CD27 | BV570 | O323 | 302825 | BIOLEGEND |
| CCR5 | BUV737 | 2D7 | 565293 | BD |
| OX40 | PECY7 | Ber-ACT35 | 350012 | BIOLEGEND |
| CD137 | APC | 4B4-1 | 309810 | BIOLEGEND |
| CD69 | PECF594 | FN50 | 562617 | BD |

\# An overlapping T-cell panel that included the tissue-resident markers CD69 and CD103 was also employed
