## Supplementary Table 1 for "Distinct Type 1 Immune Networks Underlie the Severity of Restrictive Lung Disease after COVID-19"

**Supplementary Table 1. Demographics, Co-Morbidities, Acute Treatments, and Symptoms According to Pulmonary Phenotypes.**

| **a. Demographics** | **A (*n* = 32)** | **B (*n* = 10)** | **C (*n* = 19)** | **D (*n* = 28)** | **E (*n* = 15)** | **Fisher’s Exact** |
| --- | --- | --- | --- | --- | --- | --- |
| **Sex:**  Male | 40.6% | 80.0% | 42.1% | 75.0% | 60.0% | ***P* = 0.03** |
| **Race/Ethnicity:**  White  Hispanic  Black  Other | 43.8%  34.4%  15.6%  6.3% | 30.0%  40.0%  10.0%  20.0% | 52.6%  21.1%  15.8%  10.5% | 39.3%  35.7%  21.4%  3.6% | 66.7%  26.7%  6.7%  0% | *P* = 0.36^1^ |
| **Smoking History:**  Current  Former | 0%  28.1% | 10.0%  30.0% | 5.3%  42.1% | 3.6%  32.1% | 0%  53.3% | *P* = 0.46^2^ |
| **b. Comorbidities** | **A (*n* = 32)** | **B (*n* = 10)** | **C (*n* = 19)** | **D (*n* = 28)** | **E (*n* = 15)** | **Fisher’s Exact** |
| **Diabetes:**  Type 2  Pre-Diabetes  Other^3^ | 31.3%  6.3%  3.1% (G) | 50.0%  0%  0% | 36.8%  15.8%  0% | 50.0%  3.6%  3.6% (S) | 46.7%  6.7%  6.7% (S) | *P* = 0.68^4^ |
| **Pulmonary Disease:**  All^5^  Asthma only (*n* = 27) | 37.5%  34.4% | 20.0%  20.0% | 36.8%  31.6% | 32.1%  14.3% | 53.3%  26.7% | *P* = 0.55  *P* = 0.46 |
| **Hypertension:** | 28.1% | 50.0% | 47.4% | 39.3% | 60.0% | *P* = 0.27 |
| **Hyperlipidemia:** | 25.0% | 20.0% | 36.8% | 21.4% | 6.7% | *P* = 0.36 |
| **GERD:** | 15.6% | 10.0% | 21.1% | 21.4% | 26.7% | *P* = 0.83 |
| **Sleep Apnea:**^6^ | 15.6% | 20.0% | 15.8% | 10.7% | 20.0% | *P* = 0.89 |
| **Cancer:**^7^ | 6.3% | 20.0% | 10.5% | 10.7% | 6.7% | *P* = 0.72 |
| **Anxiety/Depression:** | 15.6% | 0% | 21.1% | 10.7% | 20.0% | *P* = 0.58 |
| **Osteoarthritis:** | 9.4% | 0% | 15.8% | 10.7% | 26.7% | *P* = 0.38 |
| **Cardiac Disease:**^8^ | 9.4% | 0% | 10.5% | 7.1% | 6.7% | *P* = 0.98 |
| **Other Endocrine:** ^9^ | 6.3% | 0% | 26.3% | 3.6% | 6.7% | *P* = 0.09 |
| **Autoimmunity:**^10^ | 9.4% | 0% | 0% | 7.1% | 13.3% | *P* = 0.54 |
| **Liver Disease:** ^11^ | 6.3% | 0% | 0% | 7.1% | 13.3% | *P* = 0.55 |
| **Chronic Kidney Disease:** | 3.1% | 0% | 10.5% | 7.1% | 6.7% | *P* = 0.80 |
| **Organ Transplant:**^12^ | 0% | 0% | 0% | 10.7% | 6.7% | *P* = 0.15 |
| **c. Acute Treatment** | **A (*n* = 32)** | **B (*n* = 10)** | **C (*n* = 19)** | **D (*n* = 28)** | **E (*n* = 15)** | **Fisher’s Exact** |
| **Dexamethasone**^13^ | 46.9% | 70.0% | 73.7% | 42.9% | 73.4% | *P* = 0.09 |
| **Remdesivir**^14^ | 34.4% | 50.0% | 42.1% | 39.3% | 80.0% | *P* = 0.07 |
| **Convalescent Plasma** | 6.3% | 0% | 10.5% | 17.9% | 20.0% | *P* = 0.41 |
| **Tocilizumab** | 3.1% | 10.0% | 10.5% | 10.7% | 0% | *P* = 0.49 |
| **d. Post-COVID Symptoms** | **A (*n* = 32)** | **B (*n* = 10)** | **C (*n* = 19)** | **D (*n* = 28)** | **E (*n* = 15)** | **Fisher’s Exact** |
| **Dyspnea on Exertion** | 40.6% | 50.0% | 68.4% | 57.1% | 86.7% | ***P* = 0.04** |
| **Fatigue** | 46.9% | 60.0% | 36.8% | 32.1% | 46.7% | *P* = 0.54 |
| **Sleep Disruption** | 56.3% | 50.0% | 47.4% | 28.6% | 33.3% | *P* = 0.23 |
| **Cough** | 18.8% | 30.0% | 42.1% | 39.3% | 40.0% | *P* = 0.30 |
| **Pain/Numbness** | 34.4% | 30.0% | 42.1% | 35.7% | 40.0% | *P* = 0.96 |
| **Brain Fog/Memory Issues** | 15.6% | 20.0% | 31.6% | 14.3% | 13.3% | *P* = 0.60 |
| **Dizzy/Lightheaded** | 9.4% | 0% | 21.1% | 7.1% | 20.0% | *P* = 0.36 |

Proportion of patients within each pulmonary phenotype according to **(a)** demographics, **(b)** comorbidities at time of initial illness, **(c)** treatment received during acute COVID-19, and **(d)** patient-reported post-COVID symptoms. All entries are listed in order of prevalence within the COVID-RC. Duplicate subjects within the same cluster were removed from analysis.

^1^Comparison of White vs. All other

^2^Comparison of Any vs. Never

^3^Includes Gestational diabetes (G: *n* = 1 subject) and Steroid-induced diabetes (S: *n* = 1)

^4^Comparison of all Diabetes + Pre-Diabetes vs. None

^5^Includes TB (*n* = 3), COPD (*n* = 3), Emphysema (*n* = 3), ILD/RA-ILD (*n* = 2), IPF (*n* = 1), Sarcoidosis (*n* = 2)

^6^Includes OSA (*n* = 8)

^7^Includes Prostate (*n* = 2), Skin (Squamous cell, *n* = 1; Melanoma, *n* = 1; Basal cell carcinoma, *n* = 1), Colon (*n* = 1), Hepatocellular (*n* = 1), Abdominal tumor (*n* = 1)

^8^Includes CAD (*n* = 4), Bradycardia (*n* = 1), Angina pectoris (*n* = 1), Unspecified abnormality (*n* = 1), Atrial tachycardia (*n* = 1)

^9^Includes Hypothyroidism (*n* = 7), Hirsutism + PCOS (*n* = 1)

^10^Includes RA (*n* = 4 subjects), Sjogren’s (*n* = 1), ANA positive (*n* = 1)

^11^Includes HCV (*n* = 2), Cirrhosis (NASH, *n* = 1; Alcoholic, *n* = 1), Hepatic steatosis (*n* = 1), Gilbert (*n* = 1), Cancer (*n* = 1)

^12^Includes Liver (*n* = 3), Kidney (*n* = 2)

^13^Includes subjects who received unspecified steroids (*n* = 2)

^14^Subjects enrolled in blinded Remdesivir trial (*n* = 2) excluded from Fishers’ Exact test calculation
