## Supplementary Figures and Table of Contents for "Distinct Type 1 Immune Networks Underlie the Severity of Restrictive Lung Disease after COVID-19"

**Supplementary Figure 1.** Swimmer Plot of COVID-19 Disease Course and Follow-up Visits in the COVID-RC.

**Supplementary Figure 2.** Contextualizing the COVID-RC Within the COVID-19 Pandemic in Virginia, USA.

**Supplementary Figure 3.** Representative Chest CT Images for Subjects with Mild, Moderate, and Severe Fibrosis.

**Supplementary Figure 4.** Limited Variation in Tracking Responders Expanding (T-REX) Analysis Between Batches.

**Supplementary Figure 5.** Limited Variation in T-REX Analysis Between Uninfected Control Groups.

**Supplementary Figure 6.** Comparison of T-REX Signatures with Manually Gated T-cell Types.

**Supplementary Figure 7.** Gating Strategy Used to Identify Circulating Immune Subsets.

### Tables:

**Supplementary Table 1.** Demographics, Co-Morbidities, Acute Treatments, and Symptoms According to Pulmonary Phenotypes.

**Supplementary Table 2.** Molecular Signatures Of and Relationships Between T-cell Populations Derived from T-REX Analysis of Pulmonary Phenotypes.

**Supplementary Table 3.** Pearson Correlations of Elastic Net-selected Features and Co-correlates with All Immune Features Within Each Patient Group.

**Supplementary Table 4.** Spectral Flow Cytometry Panels.

**Supplementary Table 5.** Complete List of Immune Subsets Identified by Flow Cytometry Panel 2.

**Supplementary Table 5.** Pearson Correlations of Elastic Net-selected Features and Co-correlates with All Immune Features Within Each Patient Group.

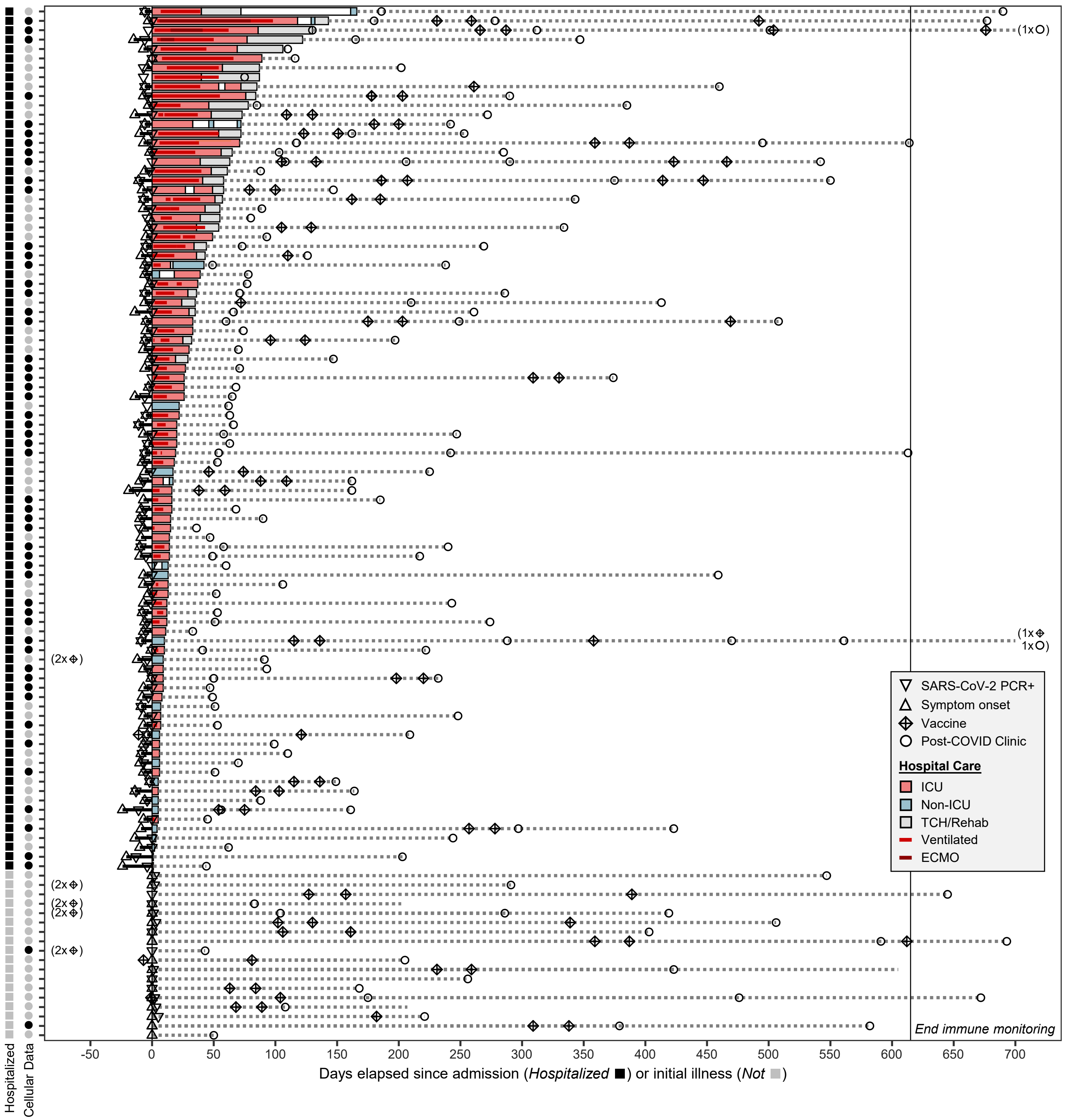


**Supplementary Figure 1. Swimmer Plot of COVID-19 Disease Course and Follow-up Visits in the COVID-RC.** Filled symbols on the left denote patients who were hospitalized (*squares*) and those for whom cellular studies were performed (*circles*).All hospitalized subjects, except one who was immune compromised, were not vaccinated at the time of initial COVID-19 illness. Four non-hospitalized subjects were vaccinated at the time of initial illness (133-458 days post-vaccination at illness). Immune data not available beyond 615 days post-COVID (vertical line).

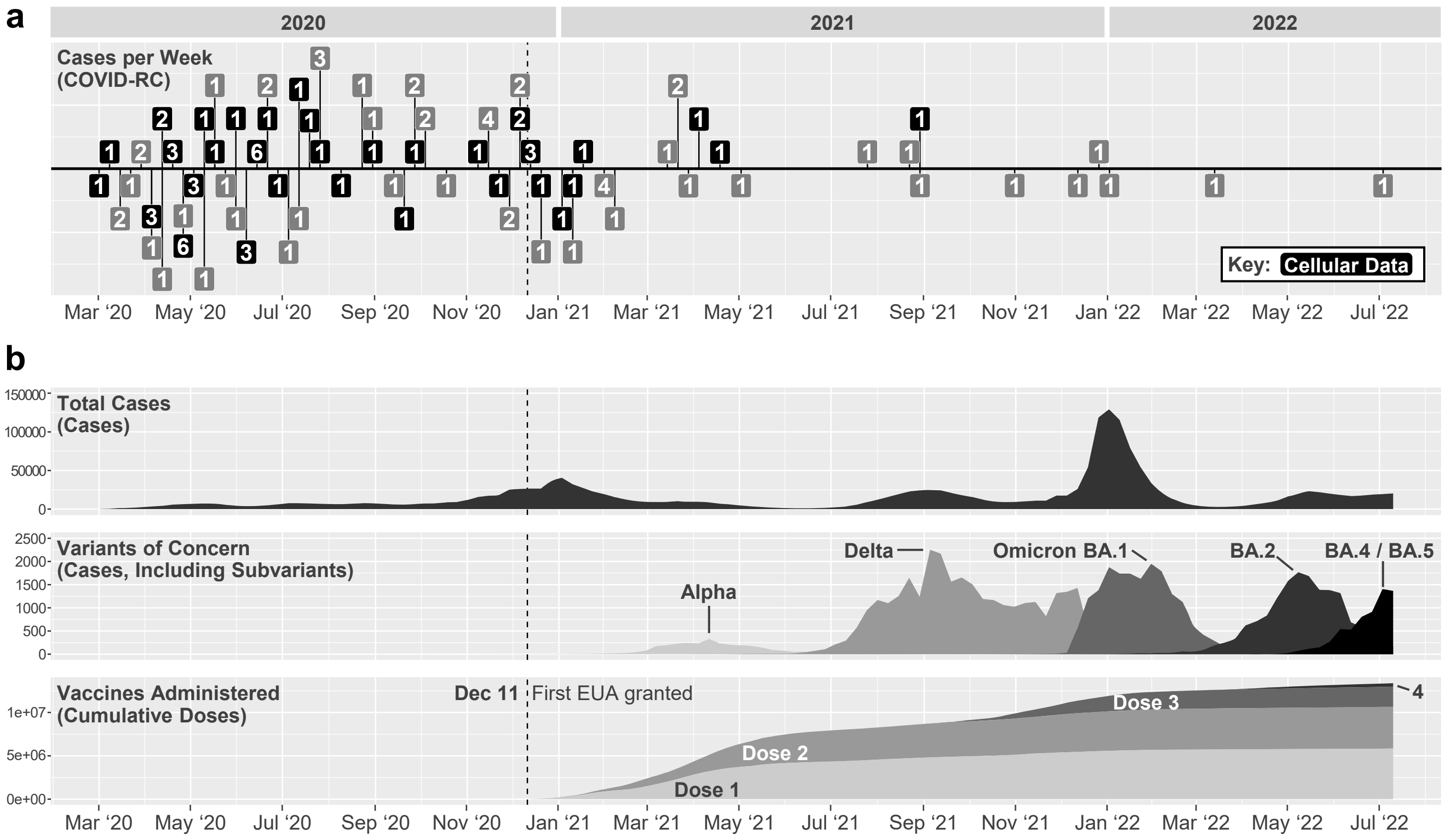


**Supplementary Figure 2. Contextualizing the COVID-RC Within the COVID-19 Pandemic in Virginia, USA.** **(a)** Number of COVID-RC subjects per week that contracted COVID-19 (earliest instance of symptoms, PCR positivity, or hospital admission). Subjects selected for cellular studies are highlighted. **(b)** Timeline of the COVID-19 pandemic in Virginia, according to weekly reports of total COVID-19 cases, variant of concern monitoring, and vaccine administration. Vertical line indicates the first COVID-19 vaccine Emergency Use Authorization (EUA). Data; Virginia Department of Health COVID-19 Public Use Datasets (data.virginia.gov)

**Mild**

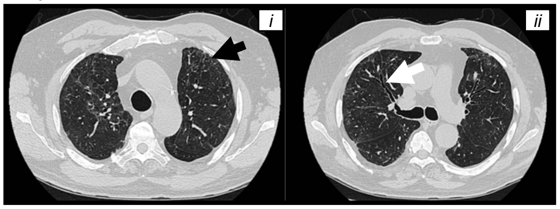


Mild persistent pulmonary parenchymal findings 9 months after COVID-19 positive test. Sub pleural reticulation (***i***, black arrow) with bronchiectasis (***ii***, white arrow) comprising no more than 25% of total lung volume.

**Moderate**

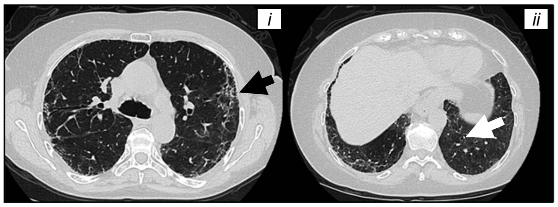


Moderate persistent pulmonary parenchymal findings 16 months after COVID-19 positive test. Small areas of peripheral honeycomb change and sub pleural banding/scar (***i***, black arrow) with moderate associated peripheral traction bronchiectasis and bronchiolectasis extending to the lung bases (***ii***, white arrow) distributed throughout 25 to 50% of the total lung volume.

**Severe**

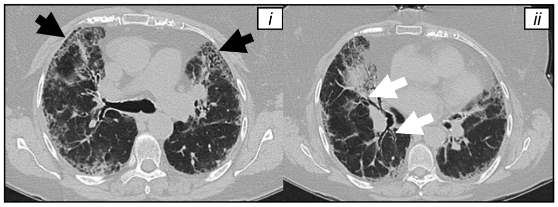


Severe persistent pulmonary parenchymal findings 9 months after COVID-19 positive test. Bilateral moderate peripheral honeycomb change (***i***, black arrows) with extensive associated peripheral traction bronchiectasis and bronchiolectasis (***ii***, white arrows) comprising more than 50% of total lung volume, including all five lung lobes.

**Supplementary Figure 3. Representative Chest CT Images for Subjects with Mild, Moderate, and Severe Fibrosis.**

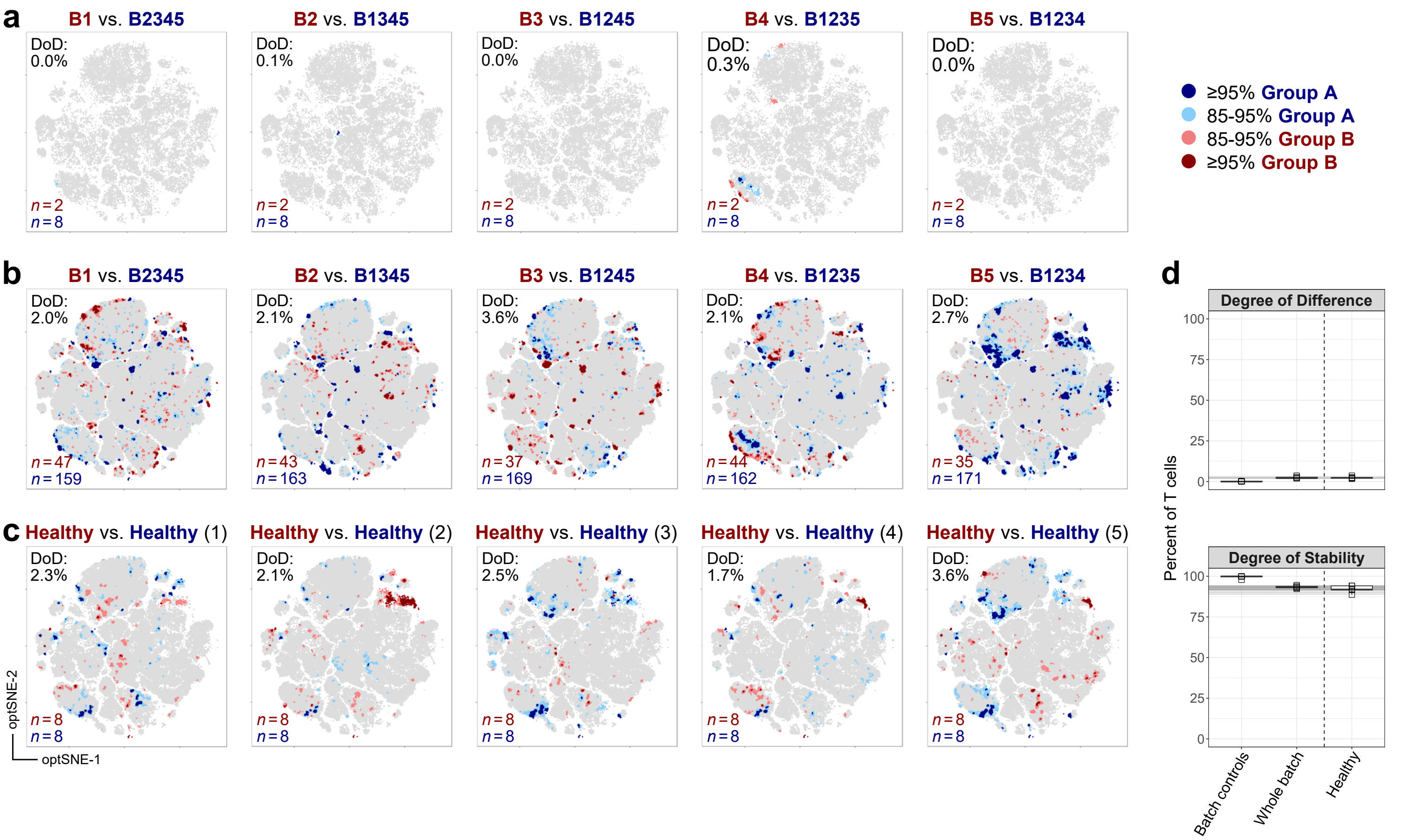


**Supplementary Figure 4. Limited Variation in Tracking Responders Expanding (T-REX) Analysis Between Batches. (a)** T-REX comparison of standard batch control samples between batches. **(b)** Comparison of all samples between batches. **(c)** Comparison of healthy control samples, iterated five times, to define variation among the healthy population. **(d)** T-REX degree of difference (DoD, percent of cells in regions of 95% enrichment) and stability (percent of cells in regions of <85% enrichment). Gray shading shows the range (*light*) and IQR (*dark*) for differences among healthy subjects, as shown in ***c***.

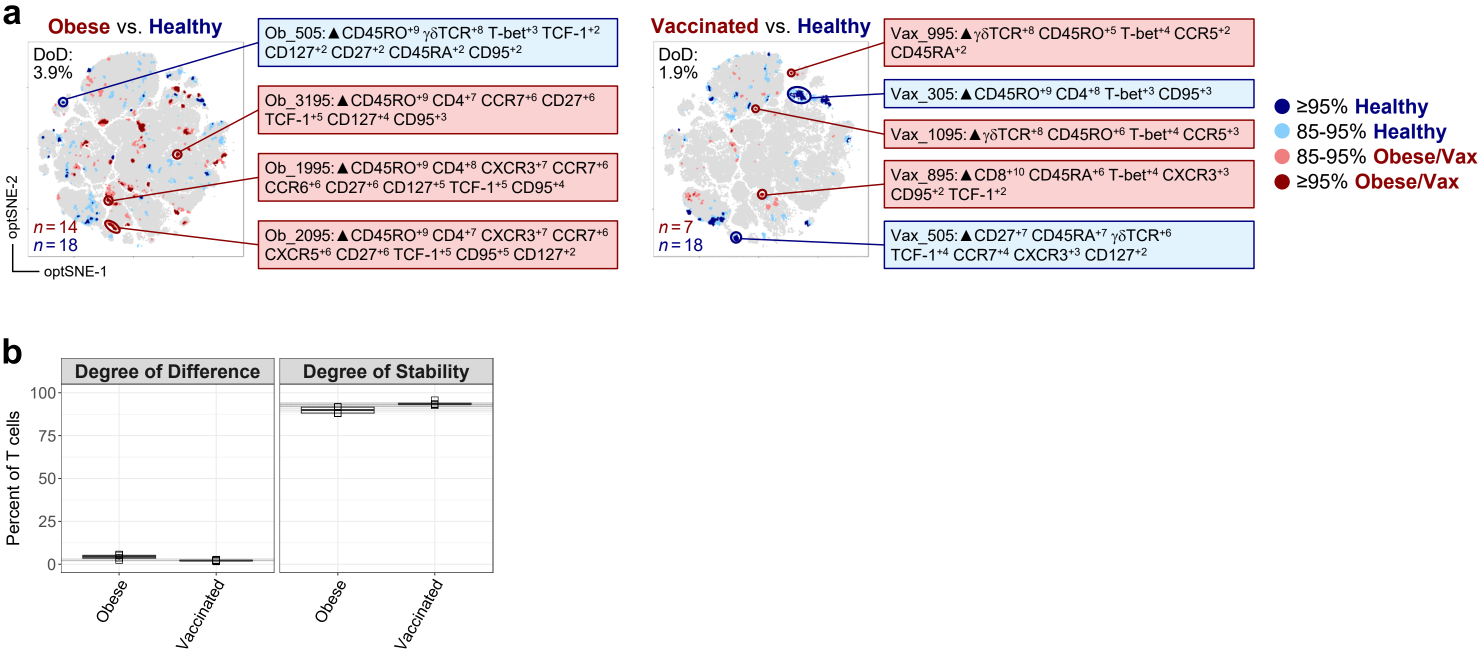


**Supplementary Figure 5. Limited Variation in T-REX Analysis Between Uninfected Control Groups. (a)** T-REX comparison of obese (*left*), and vaccinated control groups (*right*, 6 months post-vaccine) versus healthy controls (COVID-naïve). Robust populations are circled, with Marker Enrichment Modeling (MEM) labels provided in textboxes. **(b)** T-REX degree of difference (DoD, percent of cells in regions of 95% enrichment) and stability (percent of cells in regions of <85% enrichment) in obese and vaccinated groups versus healthy. Gray shading shows the range (*light*) and IQR (*dark*) for differences among healthy subjects, as shown in **Supplementary Fig. 4**.

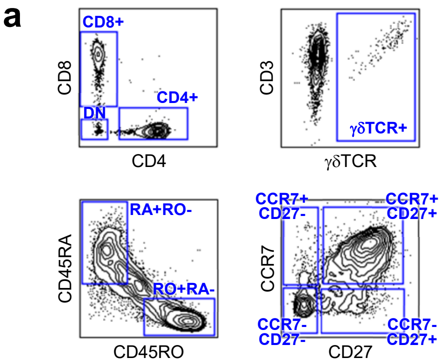


b

| Population (Manual Gating) | Boolean Filter |
|---|---|
| CD4+ T cell | **CD4+** AND NOT **γδTCR+** |
| CD8+ T cell | **CD8+** AND NOT **γδTCR+** |
| γδ T cell | **γδTCR+** |
| CD4+ γδ T cell | **γδTCR+** AND **CD4+** |
| CD8+ γδ T cell | **γδTCR+** AND **CD8+** |
| DN γδ T cell | **γδTCR+** AND **DN** |
| Naïve | **RA+RO-** AND **CCR7+CD27+** |
| $T_{CM}$ | **RO+RA-** AND **CCR7+CD27+** |
| $T_{EM}$ | **RO+RA-** AND **CCR7-CD27-** |
| TEMRA | **RA+RO-** AND **CCR7-CD27-** |

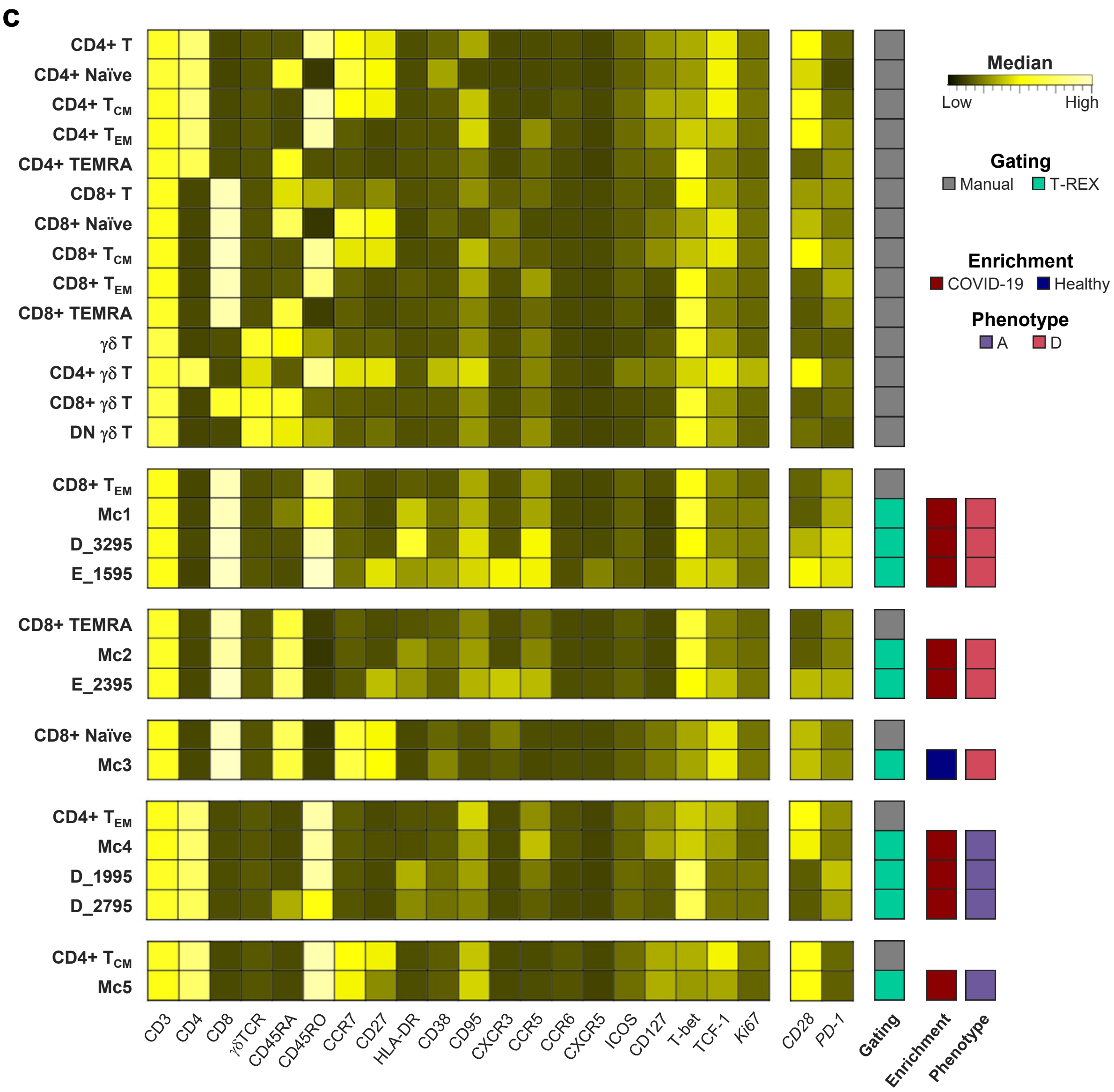


**Supplementary Figure 6. Comparison of T-REX Signatures with Manually Gated T-cell Types. (a)** Manual gates, and **(b)** Boolean gating strategy for the identification of CD4+, CD8+, and γδ T-cell subsets. All data was pre-gated for live CD3+ lymphocytes. **(c)** Median marker expression of manually gated and T-REX defined populations. For T-REX, population enrichment and the pulmonary phenotype used to identify each population are indicated. Markers in italics were not used in dimensionality reduction, T-REX, or MEM label generation.

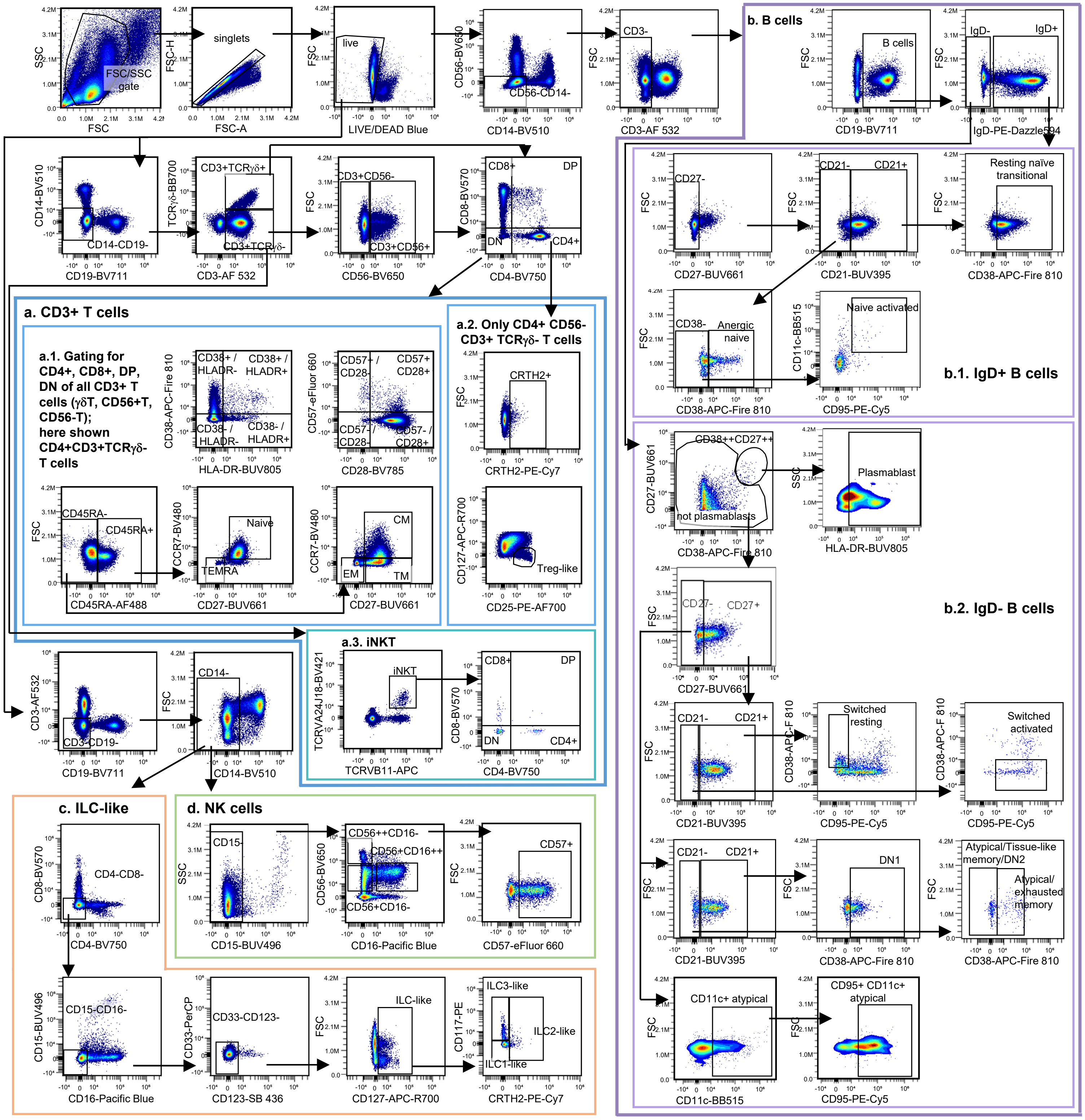


**Supplementary Figure 7.** Continued on next page.

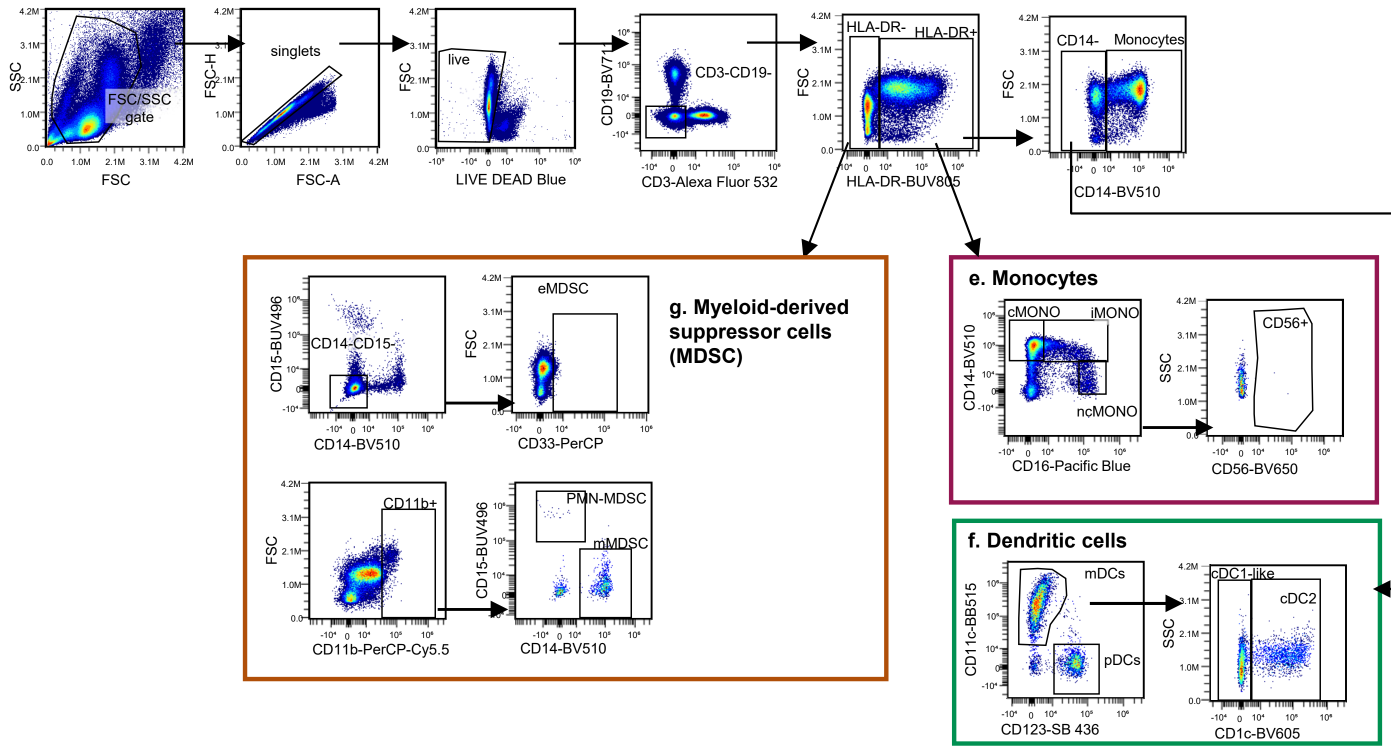


**Supplementary Figure 7. Gating Strategy Used to Identify Circulating Immune Subsets.** Gating strategy was used for the analysis of spectral flow cytometry panel 2 (**Supplementary Table 3**). Arrows indicate relationships across plots, and the gating strategy for primary populations are grouped in boxes. Data was quality checked for exact unmixing, and antibody aggregates were excluded from the analysis. Data presented is derived from frozen PBMCs of one healthy donor. Depicted gating includes the exclusion of debris, doublets and dead cells. Cells expressing lineage markers that do not define the population of interest were excluded first. **(a-blue)** T cells were defined as CD3+TCRγδ- (termed αβ) and CD3+TCRγδ+ (termed γδ). CD3+TCRγδ- were then gated as CD56+/-. All CD3+ T cell populations were further characterized based on the expression of CD4 and CD8. **(a.1)** Each CD3+ T cell population was gated for the expression of CD45RA, CCR7 and CD27 to define naïve and memory subpopulations, as well as activation markers (HLA-DR, CD38) and the senescence marker CD57 in combination with CD28. **(a.2)** CD4+ T cells were further gated for the expression of the Th2 marker CRTH2 and for markers associated to regulatory T cells ($CD127^{lo}CD25^{hi}$). **(a.3)** iNKT cells were defined as TCRγδ-CD3+ that expressed the invariant TCR (Vb11+ TCRVa24j18+), and were subsetted according to expression of CD4 and CD8. **(b-purple)** B cells were gated as CD19+ cells and divided into IgD+ (naïve) and IgD- (memory and plasmablasts) subsets. **(b.1)** Naïve (IgD+) B cells were further characterized by the expression of CD27, CD21, CD38, CD95 and CD11c. **(b.2)** Further subdivision of IgD- B cells was performed as follows: Plasmablasts were gated as $CD38^{hi}CD27^{hi}$HLA-DR+. Memory and atypical B cell subsets were gated as not-plasmablast and characterized by the expression of CD27, CD21, CD38, CD95 and CD11c. Different strategies were used for gating atypical B cells. Atypical/tissue-memory and atypical/exhausted memory subsets were gated as CD21- and distinguished by the expression of CD38 on the latter. A second strategy did not include the CD21 gate and was based on the expression of CD11c and CD95. **(c-light orange)** ILC-like cells were identified by first excluding all cells expressing lineage markers (CD3, CD19, CD14, CD15, CD8, CD4, CD16, CD33, CD123), and then by gating cells that expressed CD127. ILCs were further subdivided into ILC1 (CD117-CRTH2-), ILC2 (CRTH2+), and ILC3 (CD117+CRTH2-) subtypes. **(d-light green)** NK cells were first gated as CD3-CD19-CD14-CD15- cells. They were then classified into CD56++CD16−, CD56+CD16-, and CD56+CD16++ subsets. CD57 expression was assessed for each NK subset. **(e-magenta)** Monocytes were first gated as CD3-CD19-HLA-DR+ cells, and classified by CD14 and CD16 expression as non-classical (ncMONO; CD14−CD16++), intermediate (iMONO; CD14+CD16+), and classical (cMONO; CD14++CD16-) subsets. Expression of CD56 was assessed for each monocyte subset. **(f-dark green)** Dendritic cells (DCs) were identified first by gating on CD3-CD19-CD14-HLA-DR+ cells, and then classified by expression of CD123+ (pDCs) and CD11c+ mDCs. CD11c+ DCs were further divided into CD1c+ (cDC2) and CD1c- (cDC1-like). **(g-dark orange)** Finally, myeloid-derived suppressor cells (MDSCs) were gated as CD3-CD19-HLA-DR- cells, and then classified as early-stage (eMDSC; CD14-CD15-CD33+), monocytic (mMDSC; CD11b+CD14+CD15-) and polymorphonuclear (PMN-MDSC; CD11b+CD14-CD15+) MDSCs.
